# Bayesian inference of spike-timing dependent plasticity learning rules from single neuron recordings in humans

**DOI:** 10.1101/2023.04.20.537644

**Authors:** Ingeborg Hem, Debora Ledergerber, Claudia Battistin, Benjamin Dunn

**Affiliations:** Department of Mathematical Sciences, NTNU, Norway; Swiss Epilepsy Center, Klinik Lengg AG, Switzerland; Simula Consulting, Norway

**Keywords:** spike-timing dependent plasticity, single neuron recordings, learning rules, Bayesian statistics

## Abstract

Spike-timing dependent plasticity (STDP) learning rules are popular in both neuroscience and artificial neural networks due to their ability to capture the change in neural connections arising from the correlated activity of neurons. Recent technological advances have made large neural recordings common, substantially increasing the probability that two connected neurons are simultaneously observed, which we can use to infer functional connectivity and associated learning rules. We use a Bayesian framework and assume neural spike recordings follow a binary data model to infer the connections and their evolution over time from data using STDP rules. We test the resulting method on simulated and real data, where the real case study consists of human electrophysiological recordings. The simulated case study allows validation of the model, and the real case study shows that we are able to infer learning rules from awake human data.

## 1 Introduction

Spike-timing dependent plasticity (STDP) learning rules (Abbott & Nelson, 2000; Markram, Lübke, Frotscher, & Sakmann, 1997; Song, Miller, & Abbott, 2000) can be used to describe the evolution of neural connectivity in time and are considered to be the basis for learning and memory. They have been widely studied *in vitro* and *ex vivo* in a range of brain areas and species, including humans (Verhoog et al., 2016), and their relevance has been assessed *in vivo* (Brzosko, Mierau, & Paulsen, 2019; Jacob, Brasier, Erchova, Feldman, & Shulz, 2007; Yao & Dan, 2001). See e.g. Brzosko et al. (2019) for a review of STDP. In the STDP rules, both the sign and amount of change in connectivity depend on the relative timing of spiking neurons (Song et al., 2000), resulting in non-stationary connectivity *w*_*t*_ between neurons (Fig. 1**a**). Δ*w* is then the change in the synaptic weight when the time difference between preand postsynaptic spikes is Δ*t*. Synaptic depression will result if the postsynaptic spike occurs before the presynaptic spike (Δ*t <* 0) and vice versa for synaptic facilitation. The connection *w*_*t*_ can be constant (no change in time), stationary (random and time-independent changes), or non-stationary (time-dependent changes) (Fig. 1**b**).

**Fig. 1.**
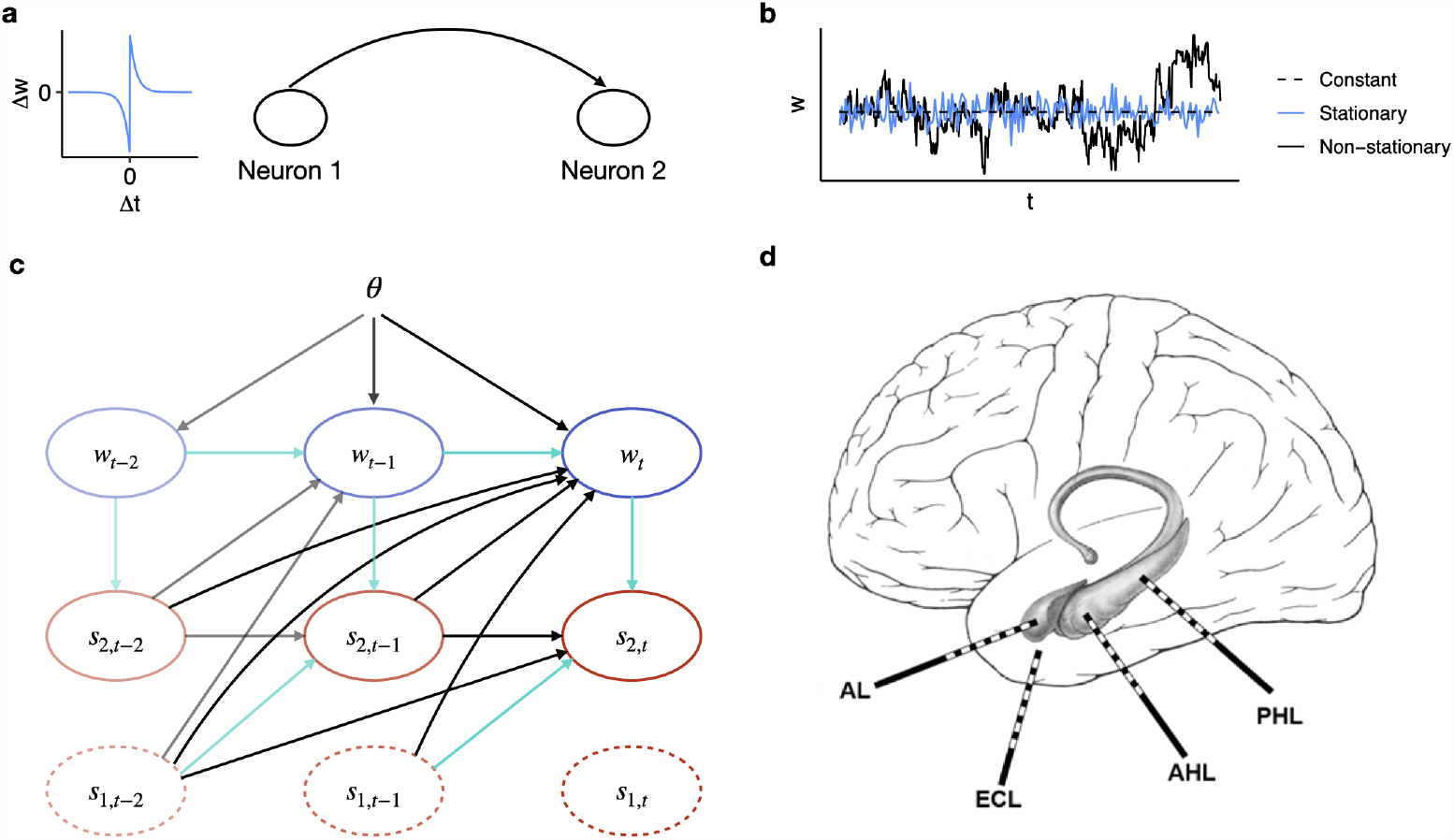
Graphical description of the learning rules and model used for inference. **a)** An example of the STDP learning rule (left) for a connection from neuron 1 to neuron 2 (right). The STDP graph shows the effect on the synaptic weight Δ*w* is the change in the synaptic weight when the time difference between preand postsynaptic spikes is Δ*t*. **b)** The connection *w*_*t*_ in time from neuron 1 to neuron 2 can be modelled as constant, stationary or non-stationary. **c)** We assume the value of the connection *w*_*t*_ at time *t* is depending on the previous connection value (*w*_*t*−1_), the previous spiking from both neurons (*s*_1,*t*_*′* and *s*_2,*t*_*′* for *t*^*′*^ *< t*) and unknown model parameters ***θ*** we want to infer. Black lines represent dependencies through the learning rule, blue lines represent dependencies through the likelihood model. **d)** We apply the model to human data recorded from the hippocampal head (AH) and body (PH), entorhinal cortex (EH) and amygdala (A). Figure shows the left (L) side of the brain.

In awake, behaving human or animal subjects it is currently not possible to assess absolute or real connectivity and plasticity between neurons (Jacob et al., 2007). By taking into account the spiking of multiple, individually recorded neurons, however, we can estimate a functional connectivity (Kobayashi et al., 2019). This can be done using generalized linear models (Pillow et al., 2008; Robinson, Berger, & Song, 2016; Roudi, Dunn, & Hertz, 2015; I. Stevenson & Koerding, 2011; Truccolo, Eden, Fellows, Donoghue, & Brown, 2005; Zaytsev, Morrison, & Deger, 2015), and prior knowledge can be incorporated by applying a Bayesian approach (Aitchison et al., 2021; Z. Chen, Putrino, Ghosh, Barbieri, & Brown, 2011; Gerwinn, Macke, & Bethge, 2010; Mishchenko, Vogelstein, & Paninski, 2011; I.H. Stevenson et al., 2009). The generalized linear models can model stationary connectivity, but more complex models are needed to infer the learning rules. Several general methods for finding plasticity rules already exist (Bengio, Bengio, & Cloutier, 1990; Jegminat, Surace, & Pfister, 2022; J. Jordan, Schmidt, Senn, & Petrovici, 2021; Lim et al., 2015; Mettler, Schmidt, Senn, Petrovici, & Jordan, 2021). The majority of these methods, however, focus on predicting the most likely rule. Linderman, Stock, and Adams (2014) suggests a way using a parametric learning rule and infer the parameters using a Bayesian framework, which allows for including prior knowledge about the learning rule. This work seeks to continue that effort.

Of particular interest is the study of learning rules in awake humans, especially in patients where a more detailed understanding of learning rule changes might provide insight into potential diseases, such as epilepsy (Bell, Lin, Seidenberg, & Hermann, 2011; Moscovitch, Cabeza, Winocur, & Nadel, 2016; Wong, 2005). Recordings from sick patients offer opportunities to study functional learning rules that could be used to generate hypothesis about the role those learning rules have in the disease. Those hypothesis could in turn for example be tested *in vitro*.

Statistical models have been used to model and infer learning rules in simulated data (S. Chen, Yang, & Lim, 2023; Linderman et al., 2014) and in experiments where memristors mimic synapses (Wu, Moon, Zhu, & Lu, 2021), but even though the technology to record spike data is available (Jun et al., 2017), not much work currently exists on how to use such recordings to infer learning rules in awake humans. We present a framework with which we aim to study synaptic plasticity in the awake human and animal brain, by assuming that the connections between pairs of neurons are driven by an STDP learning rule, and through that inferring parameters ***θ*** describing this rule. Our model is inspired by the framework presented by Linderman et al. (2014), but in a discrete approach with binary observations of neuron spikes and at a more resolved time scale. The model can be represented in a network graph describing the dependencies in the neural spike data ***s***_***i***_ and unknown synaptic weights (parameterizing the connections) ***w*** = (*w*_1_, *w*_2_, …, *w*_*T*_) for two neurons (Fig. 1**c**). We include dependencies both through the assumed likelihood model and the STDP learning rule. To uncover the unknown connections we use a Bayesian framework and assume a Bernoulli data model. We test the resulting method on both simulated data and human spike recording data from the hippocampus, entorhinal cortex and amygdala (Boran, Hilfiker, Stieglitz, Sarnthein, & Klaver, 2022) (Fig. 1**d**).

Through the simulated case study, we show that our Bayesian framework is able to infer the parameters of the STDP rule, and recover the true (simulated) values. In spiking neuron data from human intracranial recordings, we show that the model is also able to infer the parameters from real data. Our results align with experimental results previously found in animals.

## 2 Methods

We present a Bayesian framework for inferring spike-timing dependent plasticity (STDP) learning rules in spike data, that takes inspiration from Linderman et al. (2014), where spike counts are treated as observed variables, whereas dynamic, directed synaptic pairwise connections are regarded as hidden. We put the spike counts in bins, which yields binary data *s ∈ {*0, 1*}*, for both putative pre-synaptic (***s***_1_) and post-synaptic neurons (***s***_2_). We assume that the spike data for the post-synaptic neuron, *s*_2,*t*_, follows a Bernoulli process. The mean of the Bernoulli process (likelihood) is a linear combination of background noise, covariates (if any), and pre-synaptic spikes from neuron 1 at a fixed lag multiplied with a weight trajectory modelling the connection between the two neurons. This connection, denoted *w*_*t*_, is assumed to change in time and to be driven by an STDP learning rule. We use a parametric representation of the learning rule, involving parameters for maximum synaptic modification and for time intervals where synaptic strengthening and weakening occur. The goal of the modelling is to infer these parameters to get an idea of the shape of the STDP learning rule in human spike data.

### 2.1 Human spike data

To test the usability of the model on real neural data, we fit it to data from (Boran et al., 2022). This dataset contains single neuron spike trains from 13 epilepsy patients. To ensure that we study connections that likely are more than random noise, we pre-select 105 neuron pairs using temporal correlations of the spike trains (Stark & Abeles, 2009) prior to the inference (see Appendix B for details). Those 105 neuron pairs came from seven of the subjects. During this procedure we also determine which connections are excitatory, and which are inhibitory (see Methods). For each of the 105 neuron pairs we have three different sets of covariates: No covariates (cov0), covariates indicating trial difficulty (covD, four levels of difficulty based on work load in the trial), and covariates indicating trial stage (covS, four trial stages: Warning and fixation, encoding, maintenance and test). See Boran et al. (2022) for details.

The area each neuron is recorded from is given in the data, in addition to which (if any) areas are seizure onset zones for each subject. The data is recorded during a visual memory task consisting of 192 trials with four different levels of difficulty. In addition to the spike trains, we also extract information on the difficulty of each trial and when each stage of the trial occurs, as well as the pathology, gender and age of the subjects. For a given subject and session, we extract the spike times for each neuron, and create one time series with all the trials. Temporal correlation between the spike trains and a requirement of at least 5 spikes/second on average yielded 105 neuron pairs from 7 subjects (Table 1). Two subjects participated in two sessions.

**Table 1.** Spike recordings from Boran et al. (2022). Age of subjects in parenthesis. S1 and S2 represent different sessions. (S, D) denotes the number of neuron pairs in the same (S) and in two different (D) brain areas. Subject A, F and G are male. The number of connections with satisfactory high spike rates found varies between subjects, from 1 to 41. Of the 105 neuron pairs, 61 are across different parts of the brain, and 44 are within the same brain area. We only consider neuron pairs recorded with two different electrodes to ensure we consider two different neurons in each pair. 57 neuron pair connections are excitatory and 48 are inhibitory. The 105 neuron pairs are from 7 different subjects, where two of those have participated in two sessions.

| Subject | Session | Excitatory (S, D) | Inhibitory (S, D) | Total |
| --- | --- | --- | --- | --- |
| **A** (31) | S1 | (0, 1) | (0, 0) | 1 |
| **B** (56) | S1 | (0, 0) | (1, 0) | 1 |
|  | S2 | (2, 1) | (0, 0) | 3 |
| **C** (20) | S1 | (1, 0) | (3, 0) | 4 |
| **D** (51) | S1 | (2, 5) | (3, 3) | 13 |
|  | S2 | (4, 10) | (3, 3) | 20 |
| **E** (18) | S1 | (3, 0) | (3, 0) | 6 |
| **F** (47) | S1 | (4, 15) | (11, 11) | 41 |
| **G** (39) | S1 | (3, 6) | (1, 6) | 16 |
| **Total** | 9 | 19 + 38 = 57 | 25 + 23 = 48 | 105 |

### 2.2 Bayesian model for inferring learning rules

To infer the parameters of the STDP learning rule in spike data, we first consider binned spike trains ***s***_*i*_ of length *T* . Each time bin is of size 2 ms, and time bin *t* covers the events in the time interval ((*t*− 1) *·* 2ms, *t ·* 2ms] (except the first time bin which also includes 0). For a given neuron *i* and time bin *t, s*_*i,t*_ = 1 if the neuron spiked (at least once) in time bin *t*, and *s*_*i,t*_ = 0 else. We take the post-synaptic spikes as the response and build a likelihood model with the pre-synaptic spikes as input. The natural choice of likelihood model for binary spike train data ***s***_*i*_ is the Bernoulli model:

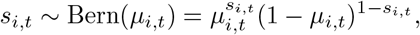

where the linear predictor *η*_*i,t*_ = logit(*μ*_*i,t*_) is given by

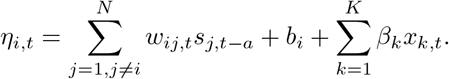

*i* and *j* indicate different neurons, *a* is the time lag for this specific connection found using the temporal correlation between the spike trains, *b*_*i*_ is an intercept representing background noise, and *β*_*k*_ is the parameter of the covariate ***x***_*k*_ = (*x*_*k*,1_, …, *x*_*k,T*_)^T^. *w*_*ij,t*_ describes the connection from neuron *j* to neuron *i* at time *t*, and *w*_*ij,t*_*?*= *w*_*ji,t*_. We model the connection, or weight trajectory, *w*_*ij,t*_ as a random walk of first order:

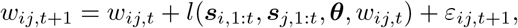

where *ε*_*ij,t*+1_ ∼ 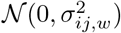 is Gaussian noise (here: synaptic noise) and ***θ*** = (*A*_+_, *A*_−_, *τ*_+_, *τ*_−_, *w*_min_, *w*_max_) are the learning rule parameters. The initial value of the weight trajectory *w*_*ij,t*_ is denoted *w*_*ij*,0_. The weight trajectory is restricted to be non-negative (*w*_*ij,t*_ ≥ 0) for an excitatory connection, and non-positive (*w*_*ij,t*_≤ 0) for an inhibitory connection. *l*(*·*) is a learning rule, and we considered a learning rule with additive dependence on the weight *w*_*ij,t*_ (additive STDP learning rule):

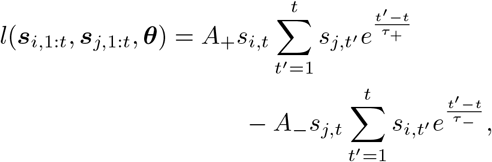

and a learning rule where the dependence is multiplicative (multiplicative STDP learning rule):

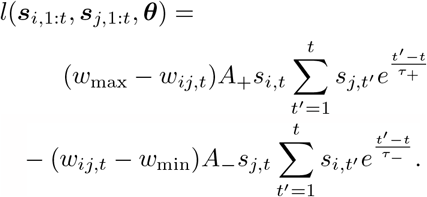

*A*_+_ and *A*_−_ represent maximum synaptic modification and *τ*_+_ and *τ*_−_ represent time intervals where synaptic strengthening and weakening occur. Song et al. (2000) investigate which parameters for the STDP learning rules fit best with experimental data from rats and other animals, and find that *A* = *A*+ = *A*_−_*/*1.05 = 0.005 and *τ* = *τ*_+_ = *τ*_−_ = 0.02 s yields simulated results close to the experimental results in frogs and rats (Bi & Poo, 1998; Debanne, Gähwiler, & Thompson, 1998; Feldman, 2000; Markram et al., 1997; Zhang, Tao, Holt, Harris, & Poo, 1998).

The goal of our analysis is to infer the learning rules for pairs of neural connections, and investigate how the whole network of neurons changes in time was outside the scope. Due to this, we consider one neuron pair at a time, and only the connection from the first neuron (neuron 1) to the second (neuron 2), such that the model simplifies to *s*_2,*t*_ ∼ Bern(*μ*_2,*t*_) and *η*_2,*t*_ = *w*_*t*_*s*_1,*t*−*a*_ + *b*_2_ + 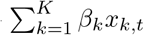 where we simplify the notation by setting *w*_21,*t*_ = *w*_*t*_. This means we assume that the only neuron affecting neuron 2 is neuron 1, and all other influence is assumed to be background noise, included in the intercept *b*_2_. We use this assumption even though one neuron may have connections to multiple other neurons, and the same neuron can be neuron 1 in one pair, and neuron 2 in another. Other neurons are assumed to lead to a systematic bias, affecting both neurons equally, which may affect the magnitude but not the shape of the weight trajectory.

Due to the complexity of the model, we employ some simplifications to reduce the computational expense during inference: *A*_+_ = *A*_−_*/*1.05 = *A, τ*_+_ = *τ*_−_ = *τ* (Song et al., 2000), *w*_min_ = 0 and *w*_max_ = 1 for excitatory connections and *w*_min_ = −1 and *w*_max_ = 0 for inhibitory connections (Morrison, Diesmann, & Gerstner, 2008). We also simplify the computation of the learning rule by only summing over the previous 10 *τ* seconds, as the contribution further back in time is essentially zero, which is also supported by experimental results (Song et al., 2000). In addition, we fix the noise parameter of the weight trajectory.

### 2.3 Inference method

The model is non-linear and not latent Gaussian due to the learning rule. This makes the model computationally expensive, but it also gives a flexible model. We use a Bayesian framework for the inference. Bayesian models require prior distributions for the model parameters *A, τ, b*_2_, *w*_*ij*,0_, and *β*_*k*_, but it also provides the opportunity to include prior knowledge about these parameters. In a hierarchical model, such as the one we consider, the difficulty of inferring a model parameter depends on the distance to the data level of the model (Goel & Degroot, 1981). Intercepts (*b*_2_) and fixed effects (*β*_*k*_) are relatively easy to infer, while the learning rule parameters and weight trajectory are more difficult. The latter is especially true for *τ*, which requires large amounts of informative data to be inferred well. This means that the intercept and fixed effects can have vague priors, while the learning rule parameters require stronger priors. Overly strong prior distributions can however yield posterior distributions highly similar to the priors, as the sampler is not able to investigate the full parameter space, so caution is needed. Based on this and the findings of Song et al. (2000), we choose the following prior distributions: *A* ∼ *π*(*A*) = Gamma(2, 200), *τ*∼ Gamma(4, 100), *w*_0_ ∼ *𝒩*_≥0_(1, 5^2^) (truncated Gaussian distribution) for excitatory connections and *w*_0_ ∼ *𝒩*_≤0_(−1, 5^2^) for inhibitory connections, *b*_2_ ∼ *𝒩* (0, 10^2^), and *β*_*k*_ ∼ *𝒩* (0, 5^2^). We fix the noise of the weight trajectory to 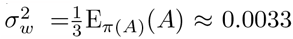to ensure that the evolution of *w*_*t*_ rarely is no more than 50% noise at time steps with spiking. E_*π*(*A*)_(*A*) is the expected value of *A* in the prior.

We use a Metropolis-Hastings (M-H) algorithm (Chib & Greenberg, 1995) to fit the model (see Appendix A for details). The background noise *b*_2_ and the parameters of the covariates *β*_*k*_ are updated independently using Gibbs steps (Casella & George, 1992), while *A, τ* and *w*_0_ are sampled together in a M-H step. In the same M-H step we also sample the weight trajectory *w*_*t*_ using a particle filter (N.J. Gordon, 1993), by precomputing the learning rule based on the current values of *A* and *τ* . To increase the efficiency of the algorithm, we use adaptive variance adjustment (Haario, Saksman, & Tamminen, 1999).

Approaches with adaptive sampling lead to correlated samples. To ensure that the algorithm has sufficient mixing of the Markov chains, we use effective sample size to compute the number of independent samples from the posterior distribution that contain the same information as our correlated samples (Plummer, Best, Cowles, & Vines, 2006).

We compare the Bayesian model to a model assuming a static (constant) weight trajectory, i.e., the maximum likelihood model, where *w*_*t*_ = *w* for all *t*. We do not include restrictions or regulations for the maximum likelihood approach, meaning the maximum likelihood has to uncover whether the connection is excitatory or inhibitory by itself. For the comparison, we use the (insample) log-likelihood value of the posterior model. That is, we compute

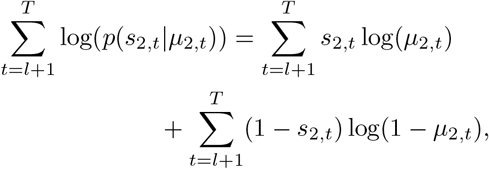

Where

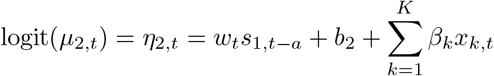

is the linear predictor with the posterior estimates. *a* is the time lag for this specific connection. For the maximum likelihood approach, we have point estimates and *w*_*t*_ = *w* for all *t*, while with the Bayesian approach, we compute the log-likelihood using each posterior sample of *b*_2_, *β*_*k*_ and the weight trajectory *w*_*t*_ and obtain a distribution for the log-likelihood. We use the median of the loglikelihood value for the comparison. In Appendix C we include simulation study results for model verification.

### 2.4 Code and reproducibility

The model is implemented in the statistical programming language R (R Core Team, 2022), and both simulated and real case studies were run on x86 64-pc-linux-gnu (64-bit) with R version 4.2.2. Necessary code for running the model is available here: https://github.com/ingebogh/synaptic_plasticity. This includes code to generate simulated data.

## 3 Results

The results are divided into two parts: We first present the main results from the simulated case study, and second the results from the real case study.

### 3.1 Model performance evaluation with simulated data

To verify that the algorithm performs as expected, we applied it to a simulated case study. We simulated data with 2 ms time bins from the true model (i.e., the model we fit to the data). Unless otherwise stated, we used the following values: *A* = 0.005 and *τ* = 0.02, *w*_0_ = 0.6 (initial value of weight trajectory), baseline firing rates for neuron 1 and 2 of 15 spikes/second (intercepts *b*_1_ = *b*_2_ = *−*3.476), *σ*_*w*_ = 0.001 (weight trajectory noise), 180 seconds of data, and an additive learning rule. We did not include covariates in these datasets, as the goal of the analysis is the learning rule parameters and not covariate parameters, but did include covariates when investigating the performance of the Bayesian model in different scenarios. We simulated 100 datasets, and to visualize the different situations that arise from the simulated, we divided them into three groups based on the evolution of the weight trajectory (Fig. 2**a**): One where the true weight trajectory approaches 0 and the connection contains little information (difference between last and first value of the weight trajectory is less than *−*0.35). One where the weight trajectory only grows and the connection (with time) becomes saturated (difference between last and first value of the weight trajectory is greater than 0.35). And one in between (difference between last and first value of the weight trajectory between −0.35 and 0.35). We denote these three groups Δ*w*_low_, Δ*w*_high_ and Δ*w*_mid_, respectively. Group Δ*w*_mid_ is the most interesting, as the two other groups are either converging to no connection and thus no learning (Δ*w*_low_) or becomes saturated with time and overfits the model in the sense that any parameter values that strengthens the connection over time will improve the model fit (Δ*w*_high_). Note that the connections does not reach saturation for the spike rates and time series lengths we use for the simulated data. The temporal correlation between spike trains method (see Section 2.1) confirmed that we can identify the correct correlation and lags (see Fig. S1 in the supplemental materials).

**Fig. 2.**
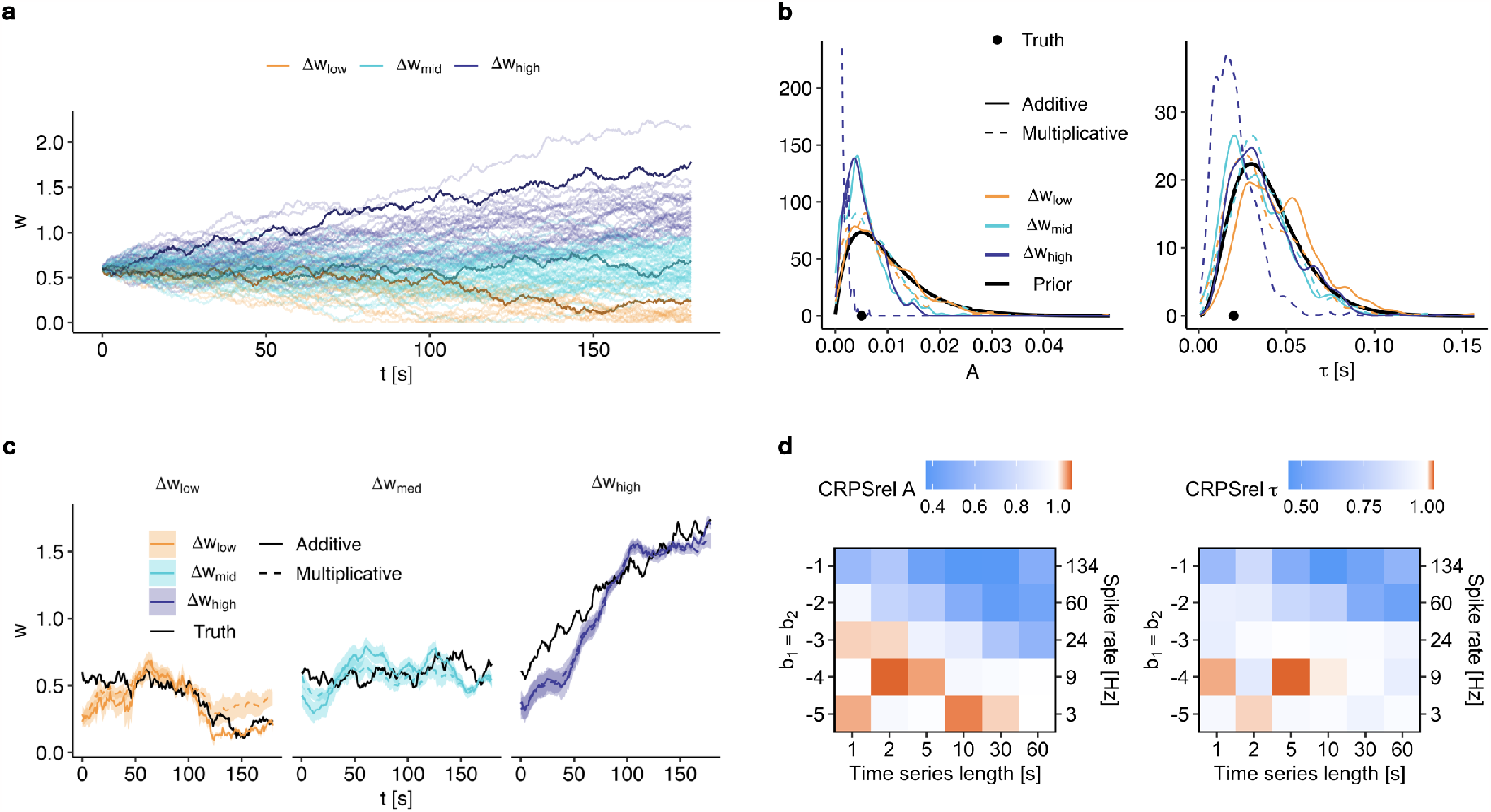
The Bayesian model uncovers the true (simulated) learning rule parameters. **a)** Weight trajectories of the 100 datasets used in the simulation study, divided into three classes Δ*w*_low_, Δ*w*_mid_ and Δ*w*_high_ (difference between last and first weight trajectory values below −0.35, between −0.35 and −0.35, and above −0.35). The results from three of the 100 datasets are displayed here, with highlighted weight trajectories in the graph. **b)** Prior and posterior distributions of *A* (left) and *τ* (right) for the three highlighted datasets. Note that the *y*-axis of the left graph is truncated at 300. For *A*, the posterior is more informative than the prior (not for dataset group Δ*w*_low_). **c)** The posterior weight trajectories *w*_*t*_*±* 1 standard deviation. The models to a large extent uncover the true (simulated) weight trajectory. **d)** The CRPSrel of *A* and *τ* for different true values of *b*_1_ = *b*_2_ and *T*, as the median over 10 simulated datasets. Only the additive learning rule is used in the model fit. CRPSrel refers to the CRPS relative to the CRPS for the prior distribution, so the prior has CRPSrel = 1. Longer spike trains with more spiking gives better values of the CRPSrel for both *A* and *τ* .

We used 2000 samples for the inference, where the first 500 are burn-in samples (sampling-based algorithms need a warm-up period) and were thus discarded. We have only included results where the effective sample size (ESS) is at least 15 (1% of 1500) for all model parameters (*A, τ, w*_0_ and *b*_2_). Early investigation of trace plots showed that the sampler has been able to explore the parameter space in a satisfactory way when ESS *>* 15. This yields 100 model fits with the additive learning rule (i.e., all converged), and 68 model fits with multiplicative learning rule. We display results from one dataset from each of the three groups, where both the additive and multiplicative models yield sufficiently large numbers of effective samples in Fig. 2**b-c**.

In the Δ*w*_mid_ group we see that the model is able to recover the true (simulated) value of *A*, and the posterior is more informative in terms of mode and uncertainty of the density, than the prior. (Fig. 2**b**). This is true for both learning rules. For *τ* we see the same trend but not as prominent. For group Δ*w*_low_, the data does not contain enough information for the model to learn about the model parameters, and the shape of the posterior does not differ much from the prior. For group Δ*w*_high_, the model with additive learning rule is able to recover the true value of *A*, while the model with multiplicative rule is not (keep in mind that the data is generated with additive learning rule). The weight trajectory *w*_*t*_ is computed at all time steps *t*, which we visualize to see how the posterior connection varies in time (Fig. 2**c**). The models with additive and multiplicative learning rules perform similar in recovering the true (simulated) trajectory *w*_*t*_. Note that any value of the parameters will increase the likelihood value when the connection *w*_*t*_ becomes saturated and keeps increasing towards infinity. However, this is not an issue for the spike rates and time series lengths we have used in the simulated datasets displayed here and we expect additional mechanisms to keep this from happening in real neurons.

When using statistical models, we must have enough information in the data to obtain reliable results. In our case, longer time series and higher spike rates of the two neurons give more informative data. We therefore study the accuracy of estimating *A* and *τ* for different values of the spike rates for the two neurons and different time series lengths (Fig. 2**d**). The accuracy is measured using the continuous ranked probability score (CRPS) (Gneiting & Raftery, 2007). The CRPS is a proper scoring rule that measures both the accuracy and the uncertainty of the estimate (a low CRPS is desired). Assessing the uncertainty in addition to accuracy is suitable in the Bayesian setting where the results are posterior distributions, and ensures that accurate but uncertain estimates are ranked worse than accurate and certain estimates. The CRPS has the mean absolute error as a special case for point estimates.

To see how the posterior distributions of *A* and *τ* compare to the prior, we report the CRPS of the parameters from the inference divided by the CRPS of the prior, which we denote CRP-Srel. The prior then has CRPSrel = 1, and values less than 1 mean the posterior is more certain about a more accurate point estimate than the prior, that is, the model has learned from the data. The median over 10 simulated datasets is used for each spike rate and time series length. We see that for higher spike rates and long time series, our model returns a posterior that is more accurate with higher certainty than the prior for both *A* and *τ*, but for low spike rates and short time series, the posterior is not performing better than the prior (Fig. 2**d**). Longer time series and more frequent spiking (higher values of *b*_1_ = *b*_2_) result in more accurate parameter estimates of *b*_2_, *w*_0_ and *w*_*t*_ (Fig. S2, left, in the supplemental materials). The times series needs to be sufficiently long for the model to lead satisfactory mixing (ESS *>* 15 for all parameters). For *w*_0_, we get a median ESS smaller than 15 for nearly half of the scenarios (combination of low *T* and low *b*_1_ = *b*_2_), indicating poor mixing of the sampler (Fig. S2, right). In longer time series with spike rates above 24 Hz, we get ESS *>* 15, meaning the sampler is able to adequately explore the parameter space.

With the simulated data, we did confirm that the Bayesian model performs as intended and that it is able to recover the true (simulated) learning rule parameters.

### 3.2 Inferring learning rules in humans

The data from Boran et al. (2022) was processed as described in 2.1, and consist of 105 neuron pairs. We fitted models with both additive and multiplicative learning rules and all covariate sets, yielding 6 model fits for each dataset. We did not use information about age, gender or brain area in the inference.

We ran the Bayesian inference algorithm for 4000 iterations for each of the 105 neuron pairs, where the first 500 are used as burn-in (warm-up) and discarded. This gives 3500 samples of each parameter and the weight trajectory at each time step. We discard all model fits where one or more of the model parameters (*b*_2_, *β*_*k*_, *A, τ, w*_0_) have effective sample size (ESS) less than 35 (as for the simulated data we used 1% of the total number of samples), as the Markov chains of those model fits did not converge. For some model fits, the sampler got stuck in parts of the parameter space and stopped due to numerical errors. These model fits were also deemed as not having converged and thus discarded. We report the number of converged models for each covariate group and learning rule type in Fig. 3**a**.

**Fig. 3.**
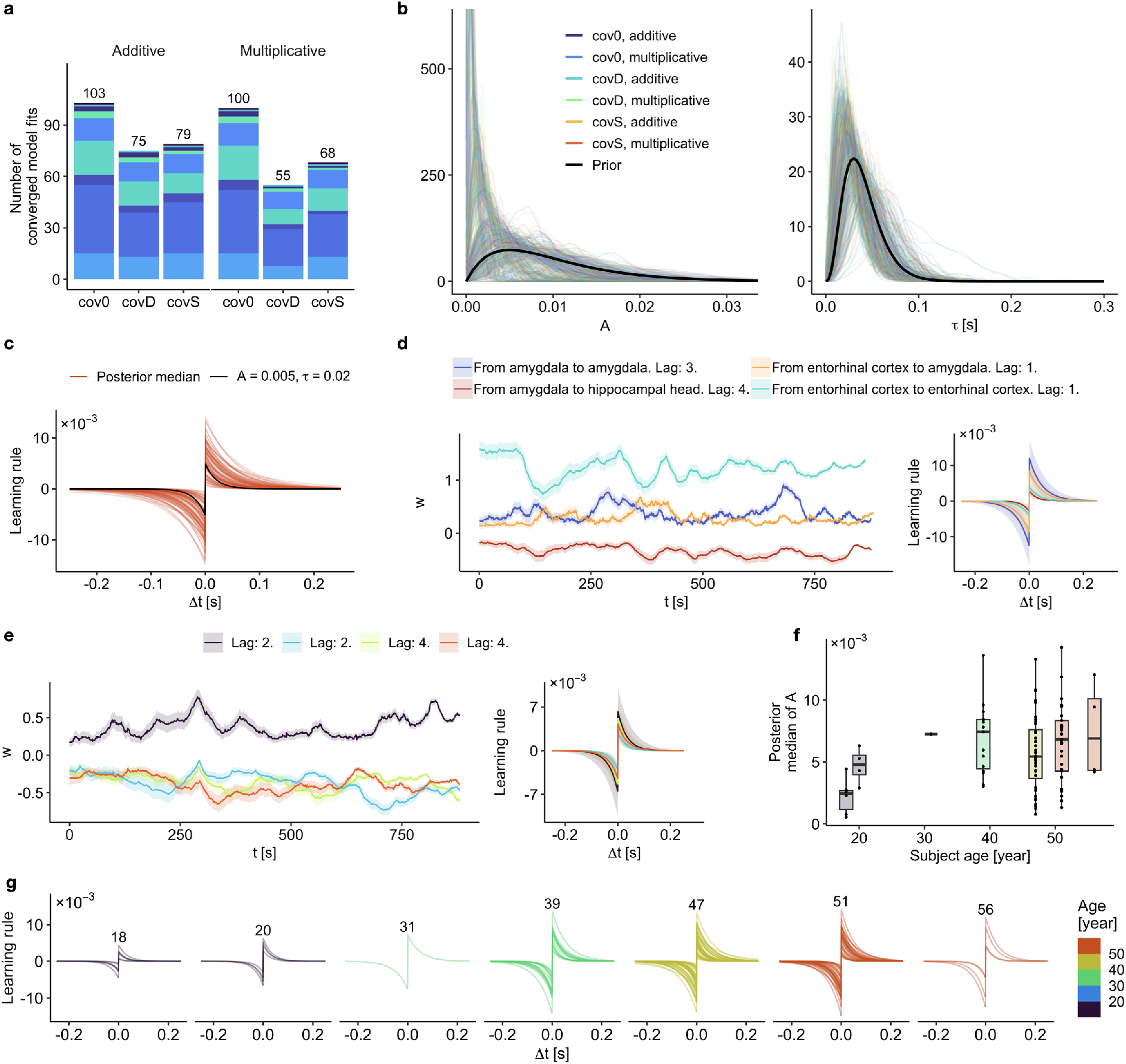
Results from the case study with neural spike trains recorded in human temporal lobes. Only results from the model with additive learning rule without covariates (cov0) are displayed in **c-g. a)** The number of converged model fits for each learning rule and covariate set, out of a total of 105 neuron pairs. Colors indicate different subjects and sessions. **b)** All posterior distributions of *A* (left) and *τ* (right), and the corresponding prior distributions. Note that the *y*-axis in the left graph is truncated at 600. **c)** Posterior median of *A* and *τ* are used to visualize the (additive) learning rule, together with the learning rule with parameters *A* = 0.005 and *τ* = 0.02. The learning rules for all 103 converged model fits are displayed in the graph. **d-e)** Posterior weight trajectories (left) and corresponding learning rules (right) from four neuron pairs. The weight trajectories are displayed with posterior mean *±* 1 standard deviation. The learning rules are based on posterior medians and corresponding uncertainties on lower and upper quartiles of *A* and *τ* . **d)** Recorded in different subjects. **e)** Recorded in the same subject and session (subject C). All neurons are located in the hippocampal head. **f)** Posterior median of the maximum synaptic modification *A*, sorted by age of subject. None of the subjects in the study were the same age. The box plots consist of the median, upper and lower quartiles, and the lowest and highest value (there were no outliers). The individual points are included as well. **g)** Posterior learning rules based on posterior median of *A* (shown specifically in **f**) and *τ*, sorted by age of subject.

With simulated data, we can evaluate the model performance by measuring how close to the truth the posterior distributions were. For real data, we cannot do this, and instead, we compared the Bayesian approach to the straight-forward maximum likelihood approach, which assumes a constant weight trajectory (*w*_*t*_ = *w*) and thus no learning rule. More specifically, for each of the 105 neuron pairs and a given covariate set, we compared the log-likelihood value of the Bayesian model (with both additive and multiplicative learning rules) to the log-likelihood value of the maximum likelihood approach as a way of assessing how well the model describes the data. For all three covariate sets, the Bayesian model (both learning rules) almost always outperformed the maximum likelihood model in terms of this loglikelihood fit. Furthermore, the Bayesian model with additive learning rule outperformed the maximum likelihood approach more often than the Bayesian model with a multiplicative rule. We obtained information on whether each connection is excitatory or inhibitory from the temporal correlation of spike trains procedure. For 9 of the neuron pairs, the maximum likelihood approach estimates an inhibitory connection (determined by the temporal correlation procedure) as excitatory. We only compare the log-likelihood for neuron pairs where the maximum likelihood estimates the correct (based on temporal correlation procedure) connection type (excitatory/inhibitory). To check that the Bayesian model behaves as we would expect in one of the cases where the maximum likelihood estimates disagree with the temporal correlation procedure, we show how the Bayesian model performs in one of those cases (Fig. S3, third panel, in the supplemental materials). We do not see anything indicating that the connection is wrongly classified using the temporal correlation: The multiplicative learning rule is small, indicating that the learning is close to non-existing, while the additive also is small compared to the other panels. However, the learning rules for one of the excitatory connection shows the same trend (Fig. S3, fourth panel). suggesting that the connection could be correctly identified with the temporal correlation.

The posterior distributions of *A* are (in general) more narrow than the prior, and the mean has shifted compared to the prior (Fig. 3**b**, left). This suggests that we have learned about the parameter from the data. The same is true for *τ* (Fig. 3**b**, right), but to a smaller extent. This is expected due to its role in the hierarchical model (Goel & Degroot, 1981). The posterior of the learning rule parameters does not seem to change between the different sets of covariates (Fig. 3**b** and Fig. S4 in the supplemental materials). In comparing the two learning rules, we see that out of the 105 neuron pairs the additive and multiplicative learning rule led to convergence in 103 and 100 neuron pairs, respectively, for the model without covariates (cov0, Fig. 3**a**). In the shared group of 100 neuron pairs where models with both learning rules converged, the additive learning rule led to the highest (best) log-likelihood value in 93 pairs, and the multiplicative led to the highest log-likelihood value in only 7. The learning rule parameters are largely not changed by the addition of covariates (Fig. S4), and also the the posterior weight trajectories *w*_*t*_ are highly similar, even though different covariate sets are used (Fig. S3). The multiplicative rule tends to yield a slightly smoother weight trajectory than the additive. More smoothing occurs when the corresponding learning rule has a small maximum synaptic modification *A*, as this results in smaller changes Δ*w*. Based on these two findings, and since our goal is to infer the learning rule parameters, we only consider the model with additive learning rule and no covariates (cov0) in the remainder of this section.

In order to compare the results from our model to previous experimental animal results, we compare the learning rules from the 103 model fits with additive learning rule and no covariates to the learning rules with parameters proposed by Song et al. (2000) (*A* = 0.005, *τ* = 0.02), which are found to match experimental results (Fig. 3**c**). We see that the posterior values from human spike recordings vary around the *A* = 0.005 and *τ* = 0.02 learning rule, indicating that the learning rules in humans resemble the ones in animals.

To visualize how the connection between two neurons vary in time, and how the connections vary between neuron pairs in the same and different subjects, we display posterior weight trajectories (Fig. 3**d**, left) and corresponding learning rules (Fig. 3**d**, right) for four neuron pairs in four different subjects. We see that the uncertainties of the learning rules are only partially overlapping between the subjects. There is however no clear pattern between the sign or magnitude of the weight trajectory and the learning rule. We also display posterior weight trajectories (Fig. 3**e**, left) and learning rules (Fig. 3**e**, right) for four neuron pairs from the same subject and session. Even though the model is fitted independently to the spike data from the four neuron pairs (the pairs have different response neurons), all four trajectories have the same increase followed by a decrease at around 300 seconds. We also see that the uncertainties in the learning rules overlap to a larger degree when all neuron pairs are from the same subject and session (Fig. 3**e**, left) than when they are from different subjects (Fig. 3**d**, left).

Identifying trends in learning rules across properties of the subjects was not the goal of our study, however, the data contains subjects of a wide range of ages (18–56), and a trend caught our attention in the process of visualizing the results. We see an increasing trend in the maximum synaptic modification *A* with increasing age (Figs 3**f** and 3**g**). However, as the number of neuron pairs analysed for each subject varies, there are more connections for high ages than for low, we cannot do any formal testing or draw any conclusions without further analysis of more data as these differences may just be coincidentally aligning with age. Note that some subjects have participated in two sessions, which is not indicated in the graphs. No trends similar to the age trend were observed when grouping the learning rules by gender, brain area, or seizure onset zones, with larger differences obvserved between subjects than between groups (Fig. S5**a-f** in the supplemental materials). Also note that the probability of discovering a spurious trend increases with the number of different groupings tested. We have only grouped the posterior learning rules based on information not used in the inference.

## 4 Discussion

We inferred the unknown, time-varying neural connections in simulated and real data using a Bayesian model. The simulated case study showed that the Bayesian approach with a binary data model works as intended, and the real case study with human brain electrophysiological recordings showed that the learning rules we found in the human brain resemble results found experimentally *in vitro* in rats and other animals.

Brain connectivity and spike-timing dependent plasticity (STDP) learning rules have long inspired the development of computer hardware and algorithms (Davies et al., 2018; Patel, Hazan, Saunders, Siegelmann, & Kozma, 2019; Vigneron & Martinet, 2020), in artificial neural networks (Diehl & Cook, 2015; Ferré, Mamalet, & Thorpe, 2018; Gilra & Gerstner, 2018; Xiang et al., 2019; Yusoff & Grüning, 2012), and in simulations of brain networks (K. Chen et al., 2021). Extended knowledge on how these learning rules work in the human brain can improve the algorithms where learning rules are employed as well as our general understanding of plasticity in the brain.

Since the data from human single neurons were recorded during a short-term memory task (Boran et al., 2022), changes in plasticity might not only be due to learning but also reflect different mnemonic items being retained in memory (Ghanbari, Malyshev, Volgushev, & Stevenson, 2017). However, to link connectivity and plasticity to a short-term mnemonic process would require orders of magnitude more data, since the item retention lasts only 3 seconds for each trial.

One of the several advantages of Bayesian inference is that we get uncertainties for our model estimates through the posterior distribution, as opposed to just a point estimate from e.g. maximum likelihood approaches. We can also easily see if the inference has utilized the data, or if the posterior is similar to the prior, the latter indicating that the model did not learn from the data. Another advantage of the Bayesian framework is the possibility of including prior knowledge. Extensive research has been done on functional connectivity and STDP learning rules (e.g., Brzosko et al., 2019; Song et al., 2000; Verhoog et al., 2016), which we can include in the model through the prior to stabilize and improve the inference. The Bayesian model outperformed the maximum likelihood approach, further supporting the idea that a constant connection is an oversimplification of the connections in the brain. Functional connectivity is not necessarily anatomical connectivity, but even though the model may be capturing something other than we intended (Das & Fiete, 2020; Dunn, Mørreaunet, & Roudi, 2015), it is still capturing something that could provide valuable insight in situations where more invasive options are not desirable. This model and method can in the future be used with the next generation of recording technology, perhaps increasing our confidence that the functional connectivity we infer is meaningful. Based on how we select neurons for the neuron pairs, the connections we investigate are not necessarily anatomical, but studying how the connections change in time and by what learning rule is still interesting and could reflect the underlying learning.

The simulated case study shows that the parameter *τ* is highly difficult to infer from the data, as the posterior does not change much from the prior. With binary data it is a difficult task for a model to infer both the weight trajectory and the learning rule parameters, and even though priors can be used to reduce the risk of overfitting, the risk is still present. More data, in our case longer time series, is one way to overcome problems with overfitting, but as more data does not necessarily mean more informative data, it may not always be a sufficient solution. To ensure that the weight trajectories do not pick up on changes in the firing rate of the neurons, a time-varying background noise effect could be included in the model. This would however further increase the complexity of the model and thus also the risk of overfitting. We recognize that there is a balance between having a model flexible enough to capture the important features and a model that is so flexible that it becomes impossible to fit properly and rapidly overfits any data provided. As our goal was to study the STDP learning rule, we have focused on a model that can recover those and at the same time keep the model as simple as possible. This is a challenging inference problem and therefore we chose a simple model with strong assumptions that allows us to still capture something that is along the lines of the STDP learning rule.

Plasticity is much more complex than what the STDP rule we use can model (Feldman, 2012; Magee & Grienberger, 2020), but as with all models, assumptions and simplifications are made in order to be able to fit the model and obtain results. A statistical model never fully describes the phenomenon of interest, but can still increase our understanding of it. The model we use is minimal, which makes it intuitive. A more biologically plausible model could, for instance, include the longer excitatory postsynaptic potential (EPSP) that our current model cannot pick up (Jegminat et al., 2022), and model membrane potential in a dynamical way instead of keeping it constant. This would, just as a time-varying background effect, lead to a more flexible but complex model at a greater risk of overfitting.

Modeling the connections between more than two neurons at the same time, or even to unobserved neurons (Battistin, Dunn, & Roudi, 2017; Dunn & Roudi, 2013), could improve the quality of the posterior distributions of the parameters, as we would have more data and thus more information to the model. This would once again increase the model complexity and potential for overfitting. Parameter sharing across pairs of neurons could increase the posterior distribution quality, but still with an increase in complexity. To allow *A*_+_ and *A*_−_, and *τ*_+_ and *τ*_−_, to be inferred independently increases the model complexity even more, but allows additional flexibility in the model and would be interesting to investigate.

A limitation with the inference method we have chosen is that it is computationally expensive, just as with all sampling-based methods, even though it converges to the true distribution in the limit. This lays restrictions on how large simulation studies and how much analysis of real data is feasible. One way to overcome this is to use approximation methods instead of, or in combination with, the sampling methods. The model we consider is close to being latent Gaussian, and for such models, a suitable method is the integrated nested Laplace approximations (INLA) method (Rue, Martino, & Chopin, 2009). By fixing the *τ* parameter of the learning rule, the model becomes latent Gaussian, and by combining sampling and INLA one could also infer *τ* (Berild, Martino, Gómez-Rubio, & Rue, 2022). The scope of this paper is however to present the Bayesian Bernoulli model, and to implement the model in a different inference framework is considered further work.

We do not expect learning to happen at all times. Further work could help to identify specific moments of learning and only including moments of learning in the inference will decrease the computational load of the model. Further work could also include investigating whether the learning rule is purely additive, purely multiplicative, a combination of the two, or even something in between additive and multiplicative (Morrison et al., 2008). Triplet-STDP learning rules could be tested with this approach (Morrison et al., 2008; Pfister & Gerstner, 2006; Yang et al., 2018), and it will be important to study if and how the learning rule parameters change over time.

In the evaluation of the learning rules, we see that there appears be a relationship between age and magnitude of the learning rule (more specifically, the maximum synaptic modification *A*) for the epilepsy patients, but we again emphasize that this may be a spurious trend, and more data and evidence are needed in order to draw conclusions. It would be interesting to investigate this further, as this is in line with a change in memory function with age (Hedden & Gabrieli, 2004), and the results from the Bayesian model give reason to formulate a hypothesis about plasticity changes with age in epilepsy patients. There are reported enhanced long-term potentiation (Kumar & Foster, 2004), reduction of learning (Orock, Logan, & Deak, 2020), and more stable synapses with aging (Mostany et al., 2013), which together appear contradictory. However, we measure functional connectivity, and one explanation for this trend is that older patients split less between different contexts such that the same neurons are active more often, in turn leading to bigger changes in the connectivity. Another explanation is that neurons in older brains lose firing precision and that their relative spike timing fluctuates more than in younger brains.

In conclusion, we show that a Bayesian framework can be successfully used to model and gain insight into the connectivity and plasticity between neurons using single neuron spiking data with known parameters (synthetic data) and from intracranial human recordings. This insight can be used to generate specific hypothesis about learning rules to be tested in future experiments.

## A Algorithms and mathematical details

In the following, we explain the details of our Metropolis-Hastings algorithm. See e.g. Chib and Greenberg (1995) for a general introduction to Metropolis-Hastings. This algorithm is an extended version of the algorithm presented in Langsrud (2020) and Myhre (2021), which again are based on the model and inference methods proposed by Linderman et al. (2014).

The overall procedure follows Algorithm 1, and consists of several sub-algorithms. With *β*_*k*_, we refer to the parameter(s) of the covariate(s), if any. For models without covariates, this parameter is not included. Ga() is a short-hand for Gamma(). *𝒩* () is the Gaussian distribution.

### Algorithm 1

Main algorithm for Bayesian inference of learning rule parameters

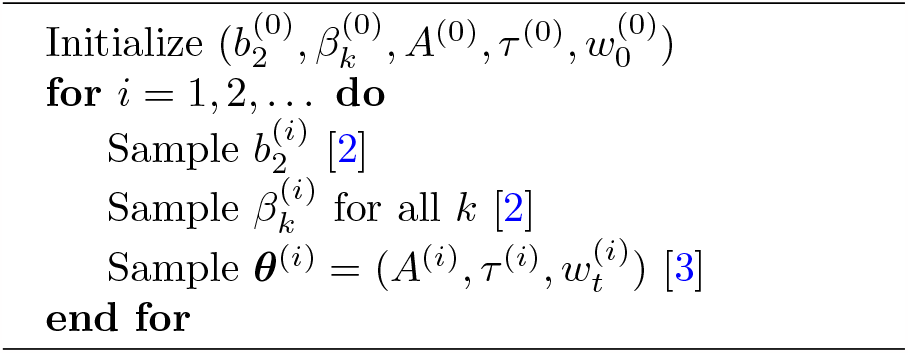

### Algorithm 2

Sample *B*^*a*^

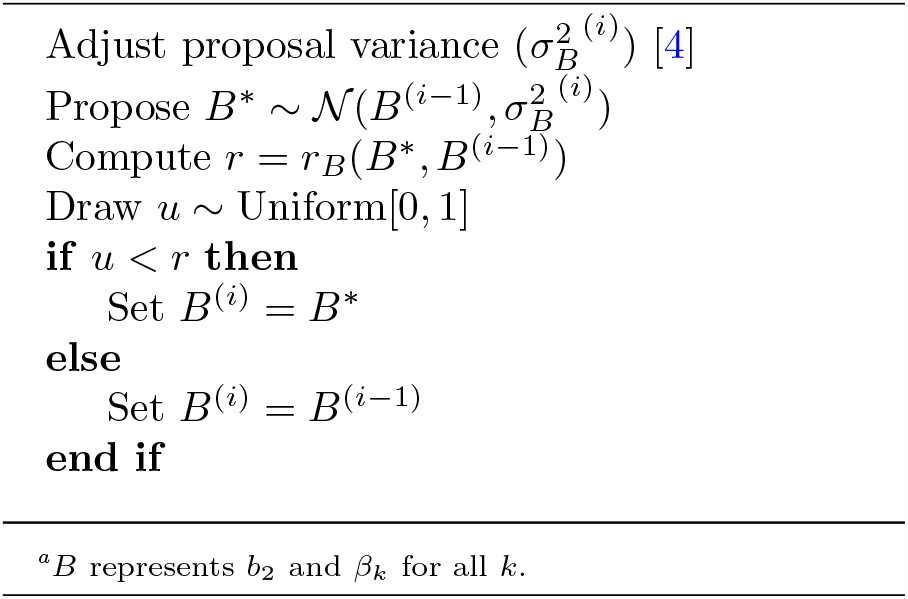

The acceptance ratio in a Metropolis-Hastings algorithm *r*_*B*_(*B*^*^, *B*^(*i*−1)^) for a parameter *B* is given by:

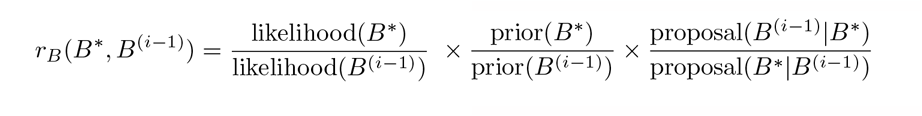

Note that the observed data ***s***, prior and proposal hyperparameters, and the model parameters not involved in that particular part of the algorithm also appear in the expression through conditioning, but are omitted here for simplicity. Also note that when the proposal distribution is symmetric around the mean, the last term is equal to 1.

### Algorithm 3

Sample *A, τ* and *w*_*t*_

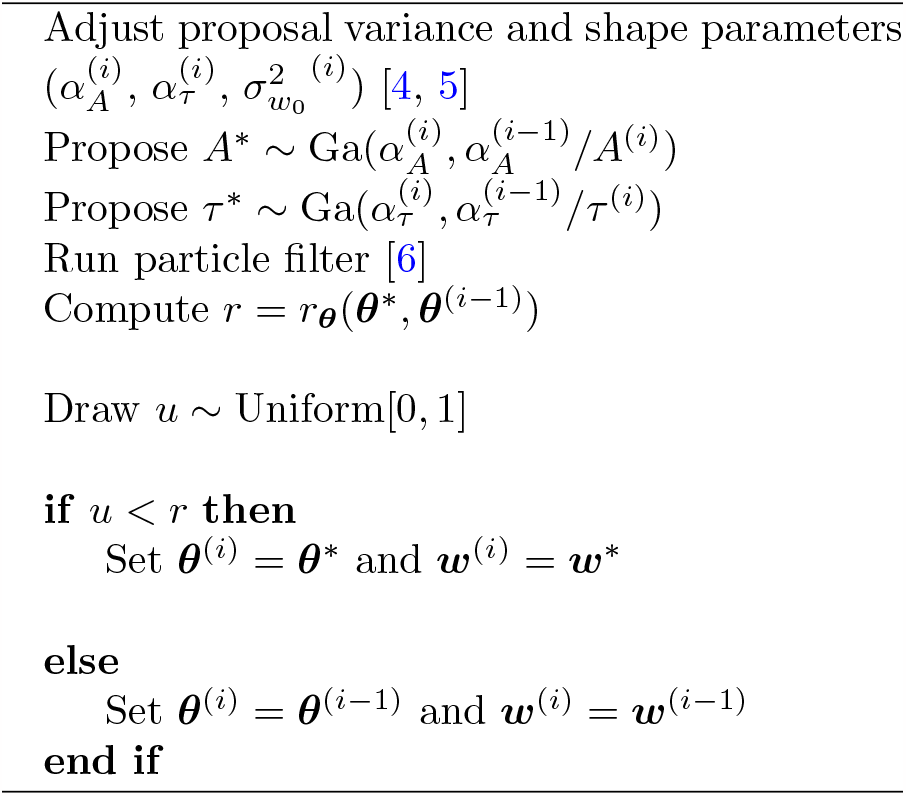

### Algorithm 4

Adjust proposal variance for parameter *ϕ*, Gaussian distribution

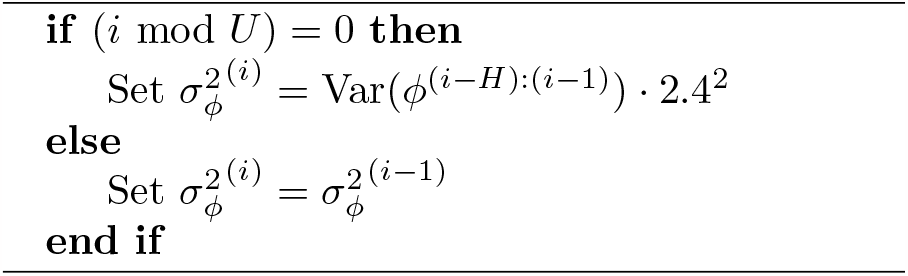

### Algorithm 5

Adjust proposal variance for parameter *ϕ*, gamma distribution

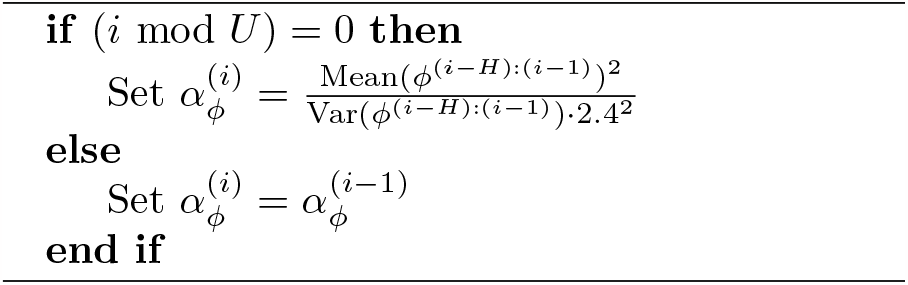

The acceptance ratio for ***θ*** is computed in a similar matter as for the parameter *B*, but since the weight trajectory *w*_*t*_ is unknown, we use a particle filter to estimate the trajectory and evaluate the likelihood (Algorithm 6).

### Algorithm 6

Particle filter

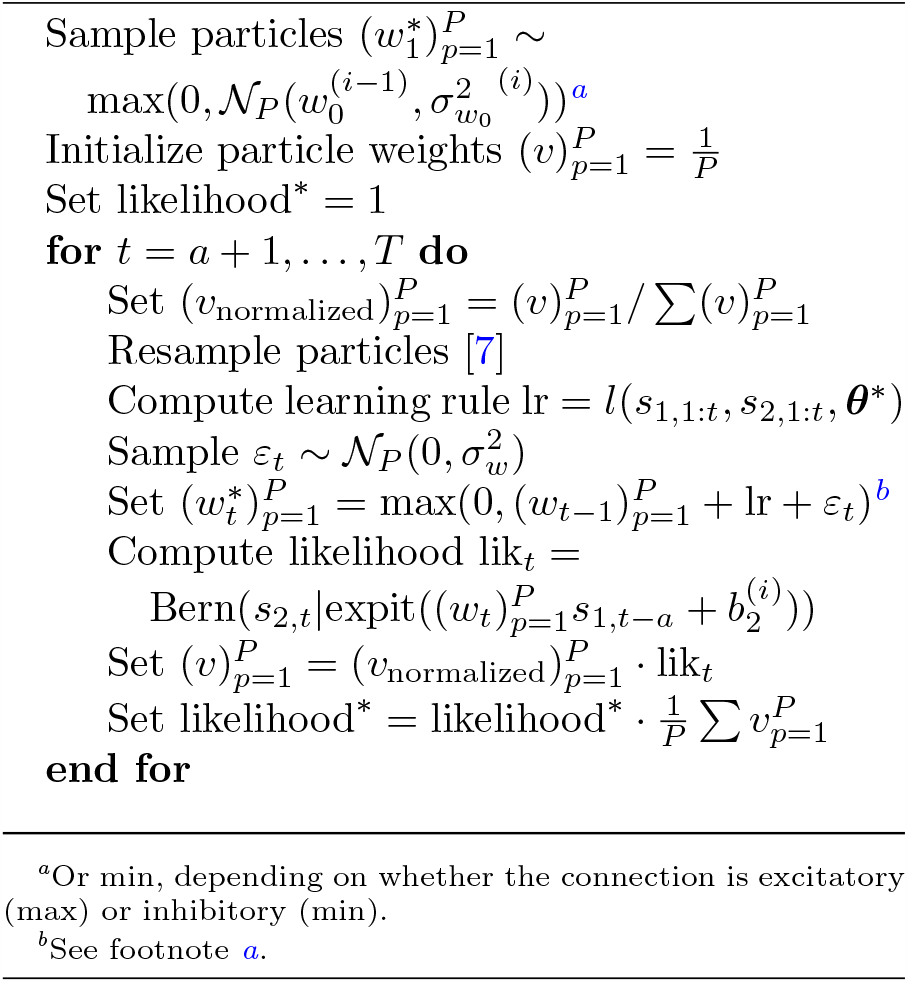

For each time step, *w*_*t*_ is given by

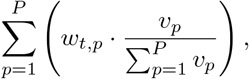

which is the weighted mean of the particles for each time step. *v*_*p*_ are the particle weights after the last time step. This is the weight trajectory used to compute the likelihood when sampling *b*_2_ and *β*_*k*_ for all *k*.

Some of the particles will have a low probability due to the random component (*ε*_*t*_). To concentrate the particles around the most probable values and avoid degeneration of the particle filter ()hmm, we resample the particles whenever the variation in the particle weights become too big (Algorithm 7). We measure this variation using the perplexity of the particle weights, such that we resample whenever

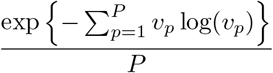

become smaller than some threshold. Martino, Elvira, and Louzada (2017) suggests to set this to about 0.66. In practice, we use 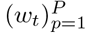 and *v*_*p*_ to compute the average *w*_*t*_ for each timestep.

### Algorithm 7

Resampling particles

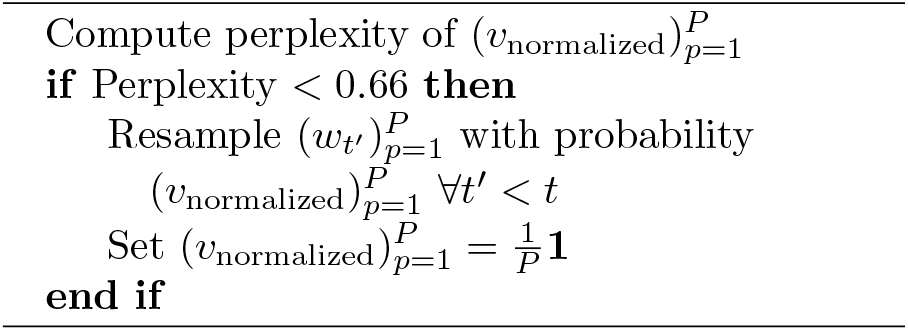

After obtaining the likelihood from the particle filter the ratio *r*_***θ***_(***θ***^*^, ***θ***^(*i*−1)^) is used to determine whether the sample of ***θ*** is accepted or not:

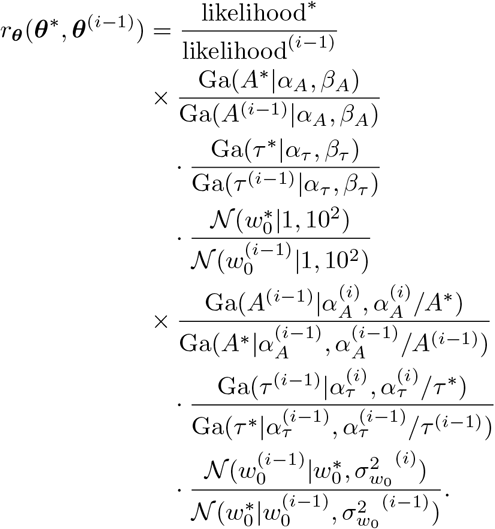

Since a neuron is either excitatory or inhibitory, we restrict the weight trajectory to have the same sign for the whole time series. To ensure this, we use a truncated Gaussian distribution for *w*_0_. To increase the speed of the algorithm, we do not do this for the weight trajectory, and instead set all negative values of *w*_*t*_ to 0, for excitatory connections, or positive values to 0, for inhibitory connections.

The prior distribution is a central component in a Bayesian framework. A good prior can yield more computationally efficient inference, and can contain prior knowledge about the problem at hand (Fuglstad, Hem, Knight, Rue, & Riebler, 2020; Gelman et al., 2020). It is however not always straight-forward to select prior distributions (Gelman, Simpson, & Betancourt, 2017; Lambert, Sutton, Burton, Abrams, & Jones, 2005).

The proposal distribution is also crucial. We use the prior distribution families as the proposals, but with different hyperparameters to increase the acceptance rate of the inference (Table 2). For all proposal distributions, we use the previous sample as the mean value, and adjust the variance according to the procedure in Haario et al. (1999) to increase the number of samples that are accepted.

**Table 2.** Algorithm settings and prior distributions. 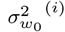is restricted to be ≥ 0.1 to ensure that the Markov chains don’t get stuck. *𝒩* (*μ, σ*^2^) is the Gaussian distribution with expected value *μ* and variance *σ*^2^. *𝒩*_≥0_() is the truncated Gaussian distribution, truncated at zero to only positive numbers.

| Parameter | Value | Prior distribution | Proposal distribution |
| --- | --- | --- | --- |
| $P$ | 200 | - | - |
| $A$ | - | $A \sim \text{Gamma}(2, 200)$ | $A^{(i)} \sim \text{Gamma}(\alpha_A^{(i)}, \alpha_A^{(i)} / A^{(i-1)})$ |
| $\tau$ | - | $\tau \sim \text{Gamma}(4, 100)$ | $\tau^{(i)} \sim \text{Gamma}(\alpha_\tau^{(i)}, \alpha_\tau^{(i)} / \tau^{(i-1)})$ |
| $w_0, w_t > 0$ | - | $w_0 \sim \mathcal{N}_{\geq 0}(1, 5^2)$ | $w_0^{(i)} \sim \mathcal{N}_{\geq 0}(w_0^{(i-1)}, \sigma_{w_0}^2{}^{(i)})$ |
| $w_0, w_t < 0$ | - | $w_0 \sim \mathcal{N}_{\leq 0}(-1, 5^2)$ | $w_0^{(i)} \sim \mathcal{N}_{\leq 0}(w_0^{(i-1)}, \sigma_{w_0}^2{}^{(i)})$ |
| $b_2$ | - | $b_2 \sim \mathcal{N}(0, 10^2)$ | $b_2^{(i)} \sim \mathcal{N}(b_2^{(i-1)}, \sigma_{b_2}^2{}^{(i)})$ |
| $\beta_k$ | - | $\beta_k \sim \mathcal{N}(0, 5^2)$ | $\beta_k^{(i)} \sim \mathcal{N}(\beta_k^{(i-1)}, \sigma_{\beta_k}^2{}^{(i)})$ |
| $\sigma_w$ | $\frac{1}{3} \mathbb{E}_{\pi(A)}(A) \approx 0.0033$ | - | - |

### B Temporal correlation between spike trains

There are several thousand neuron pairs available in the human spike recordings. The Bayesian model is too computationally expensive to fit to all neuron pairs, so we pre-select connections where we are more likely to see learning. We only consider neuron pairs where both neurons have an average spike rate of at least 5 spikes/second to ensure the data contains enough information for the analysis. To identity temporal connections in the data, we use the method described in Stark and Abeles (2009). A summary of the method and the specific settings we choose follows.

For spike trains from two neurons, we make the continuous time series into binned 0/1 time series, where for each time bin one or more spikes in a time bin interval codes to 1 and else to 0. We use a bin size of 2 ms. We keep one of the neuron spike trains fixed, and compute the number of simultaneous spikes when we shift the other spike train from *−*100 to +100 time bin lags. To avoid a bias towards time lag 0 due to a lower number of time bins to compare for high lags, we cut the ends of the spike trains such that we use the same number of time bins in the comparison for all time lags. To ensure the assumptions of further tests will not be violated, we removed all spikes but the first when the interval between two spikes is less than 6 ms (this is only done when computing the correlations, the original spike trains are used in the inference). After computing the number of simultaneous spikes for the 201 time lags (*−*100 to +100), we obtain a raw cross-correlation histogram (raw CCH).

The second step is to convolve the raw CCH with a partially hollow rectangle with a width of 11 time bins. The window hollow fraction is set to 0.42, as suggested in Stark and Abeles (2009). To prevent edge effects, bins two to six are duplicated, reversed and appended to the beginning of the raw CCH, and corresponding to the end of the CCH. This yields a predictor CCH.

Third, we compute the probability of observing the count *n* we observed or higher given the predictor CCH for each time lag *a*. We apply a continuity correction to obtain an unbiased test, and compute two tail probabilities:

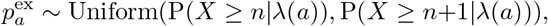

and

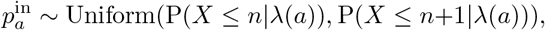

where *X* is Poisson distributed with rate *λ*(*a*) for time lag *a*, and *λ* is obtained from the predictor CCH. The first tail probability, 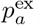, is the probability for excitatory connections (how unlikely it is that the neurons had this many simultaneous spikes, meaning spikes from the first neuron increases the probability for the second to spike), and the second, 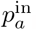, is for inhibitory (how unlikely it is that the neurons had this few simultaneous spikes, meaning spikes from the first neuron decreases the probability for the second to spike). For one neuron pair, we get these two tail probabilities for each time lag *a*. We are interested in the lags where we get small tail probabilities, and say that neurons and lags with tail probability less than 0.025 create a connection we want to analyze. Note that this method is not a formal test, and is merely used to reduce the amount of neuron pairs we study. Small tail probabilities indicate that the observed spike rate is unusual and may be caused by more than only random noise.

Since our goal is to estimate the learning rule parameters, we do not consider the zero-lag and we only consider lags up to *±* 5, corresponding to 10 ms, to capture the desired learning effects. For each neuron pair, we use at most one connection with positive lag and one with negative lag (the lag with the smallest tail probability is chosen). Just as (Boran et al., 2022) we consider the neurons recorded in different sessions with the same subject to be independent and do not connect neurons from different sessions in any way. Note that even though we consider the neuron pairs resulting from this method to form a connection, the connections may for some pairs be the result of outer factors such as common unobserved input, and we are not guaranteed that we find anatomic connections. Either way, the obtained neuron pairs show that we are able to use electrophysiological recordings to investigate learning rules in the human brain with the Bayesian model.

## C. Algorithm verification

### C.1 Model setting choices

We performed simulation studies to test the properties of the model and algorithm. We used the datasets displayed in the results from simulated data (Fig. 2A) unless otherwise stated, that is, *A* = 0.005 and *τ* = 0.02, *w*_0_ = 0.6, *b*_1_ = *b*_2_ = −3.476, *σ*_*w*_ = 0.001, 180 seconds of data, and an additive learning rule. These data are divided into three groups based on the difference of the first and last value of the weight trajectory (*w*_*T*_ − *w*_1_). The groups are Δ*w*_low_ (difference smaller than −0.35), Δ*w*_mid_ (difference between −0.35 and 0.35) and Δ*w*_high_ (difference larger than 0.35). We used time lag 1 in both the simulated data and the model. We fitted the model with both the additive learning rule and the multiplicative learning rule.

### C.1.1 Performance assessment

To assess the performance of the model, we evaluated the following: i) Continuous rank probability score (CRPS) (Gneiting & Raftery, 2007) of the model parameters, compared to the (simulated) truth. ii) CRPS of the weight trajectory, compared to the (simulated) truth, where we take the average of the CRPS over the time steps. Low values are desired. We used the R-package scoringRules (A. Jordan, Krüger, & Lerch, 2019) to compute the CRPS. iii) Effective sample size (ESS). Samples obtained from a Markov chain sampler are correlated. The ESS is a number reflecting how many independent samples the correlated samples we have correspond to. This is lower than the number of samples, and gives an indication on how well the chains have mixed during inference. High values are desired. We computed the ESS using the R-package coda (Plummer et al., 2006). We included all model fits that converged, also those with a low ESS, and report the average CRPS and ESS. In some cases, the algorithm diverged into a part of the parameter space where the parameters are so close to zero that it experiences numerical errors. The number of model fits in which this occurred are indicated in the relevant figure.

### C.1.2 Number of particles *P*

We tested the difference in performance when we use *P* = 1, 10, 50, 100, 200, 500 and 1000 particles in the particle filter in the algorithm. (Fig. 4). We display results for the different values of *P*, and the different dataset groups (Δ*w*_low_, Δ*w*_mid_ and Δ*w*_high_).

**Fig. 4.**
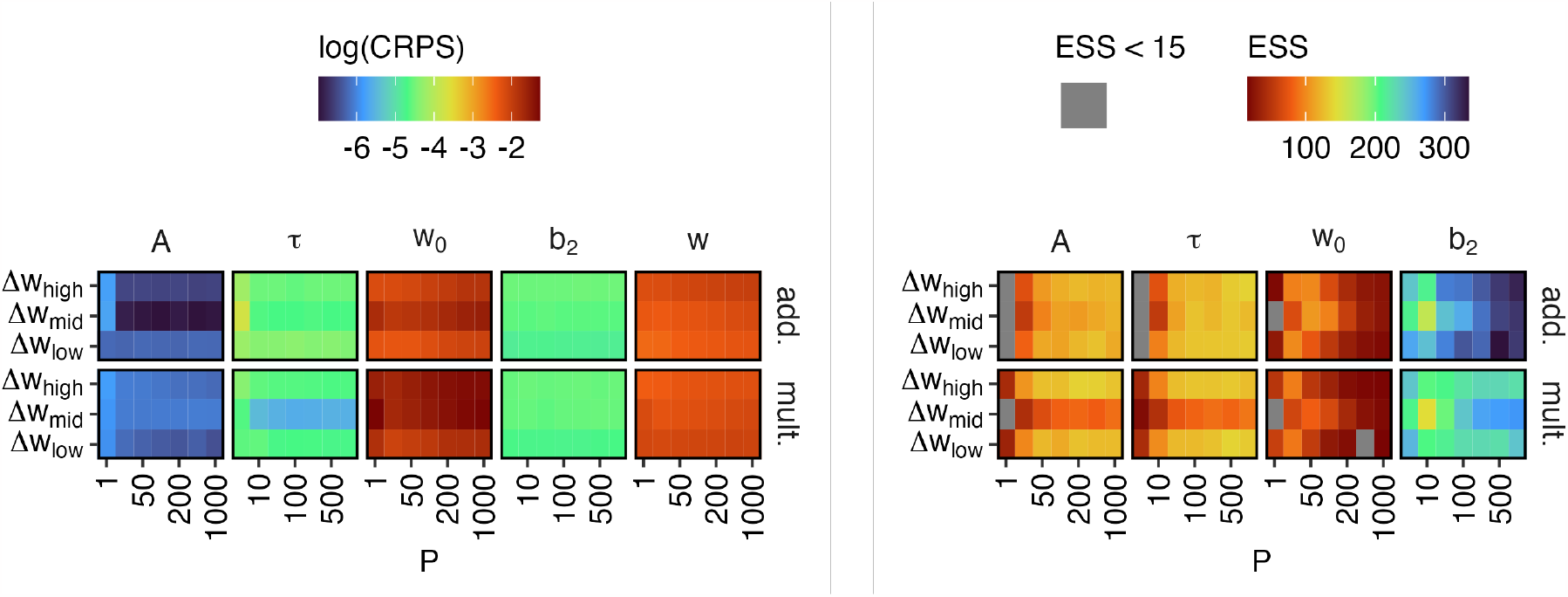
Performance for different number of particles *P* . log(CRPS) for the model parameters and weight trajectory (left) and ESS for the model parameters (right), as the mean over the 15, 53 and 32 model fits for each weight trajectory group (Δ*w*_low_, Δ*w*_mid_ and Δ*w*_high_) and value of *P* . *w* represents the whole weight trajectory. We compute the average of the CRPS, and use the logarithm when visualizing the results. Inference for 1 multiplicative model with *P* = 10 diverged.

We see that the CRPS of *A* and *τ* plateaus at around 100 particles, while the ESS stops increasing at 200 particles. The CRPS for *b*_2_ is almost constant, while the ESS for *b*_2_ increases up to 500 particles. The model with additive learning rule in general outperforms the multiplicative (this is expected since the data is generated using the additive rule). The results for the weight trajectory *w*_*t*_ and the initial value and *w*_0_ are best for *P* = 50 and becomes worse for more particles. Our focus is on the learning rule parameters *A* and *τ*, but we also need to make sure the other model parameters and effects obtain sufficient mixing in the algorithm, so we choose *P* = 200 particles in the particle filter where all parameters and effects result in good inference. The results differ only slightly between the dataset groups.

### C.1.3 Prior distributions for *A* and *τ*

We tested four different priors for *A*, and four for *τ* (Fig. 5). The prior choice for *A* mainly affects the posterior diagnostics for *A* (Fig. 6), but the other parameters are to some degree influenced as well. The Gamma(2, 400)-prior yields the best results, but has a mean equal to the true value and is deemed too strong and not optimal for new data where we do not know the truth. The Gamma(4, 50)-prior is too wide and does not lead to accurate parameter estimates, and Gamma(1, 50) shrinks the parameter to 0 and leads to poor mixing (low ESS). We therefore choose the Gamma(2, 200)-prior, as it is not too wide and not too narrow. The multiplicative learning rule model is outperformed by the additive model. The prior choice for *τ* mainly affects *τ* (Fig. 7). The Gamma(1, 100)-prior lead to lower ESS, while Gamma(2, 100) has mean equal to the true (simulated) value and is too strong. Gamma(4, 100) leads to a slightly lower and thus better CRPS for *τ* than Gamma(5, 100), so we choose (4, 100) as the prior parameters for *τ* . Again, the multiplicative learning rule model is outperformed by the additive. The weight trajectory in the most realistic dataset group (Δ*w*_mid_) is more difficult to estimate than the others, based on the CRPS value of the weight trajectory (Figs 6 and 7).

**Fig. 5.**
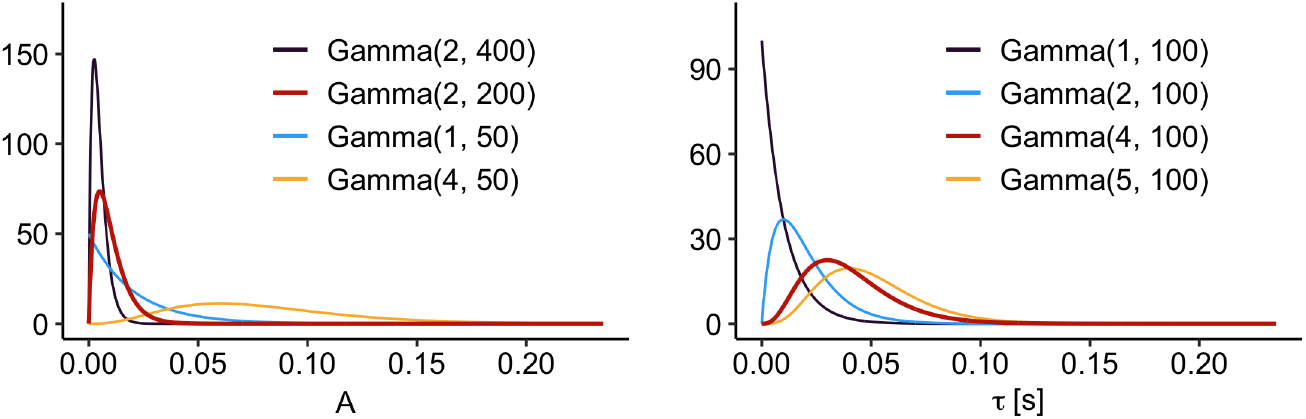
Prior distributions for learning rule parameters. The prior distributions for *A* (left) and *τ* (right) investigated in the simulation study. The bold priors are the chosen ones (*A* ∼ Gamma(2, 200) and *τ* ∼ Gamma(4, 100)).

**Fig. 6.**
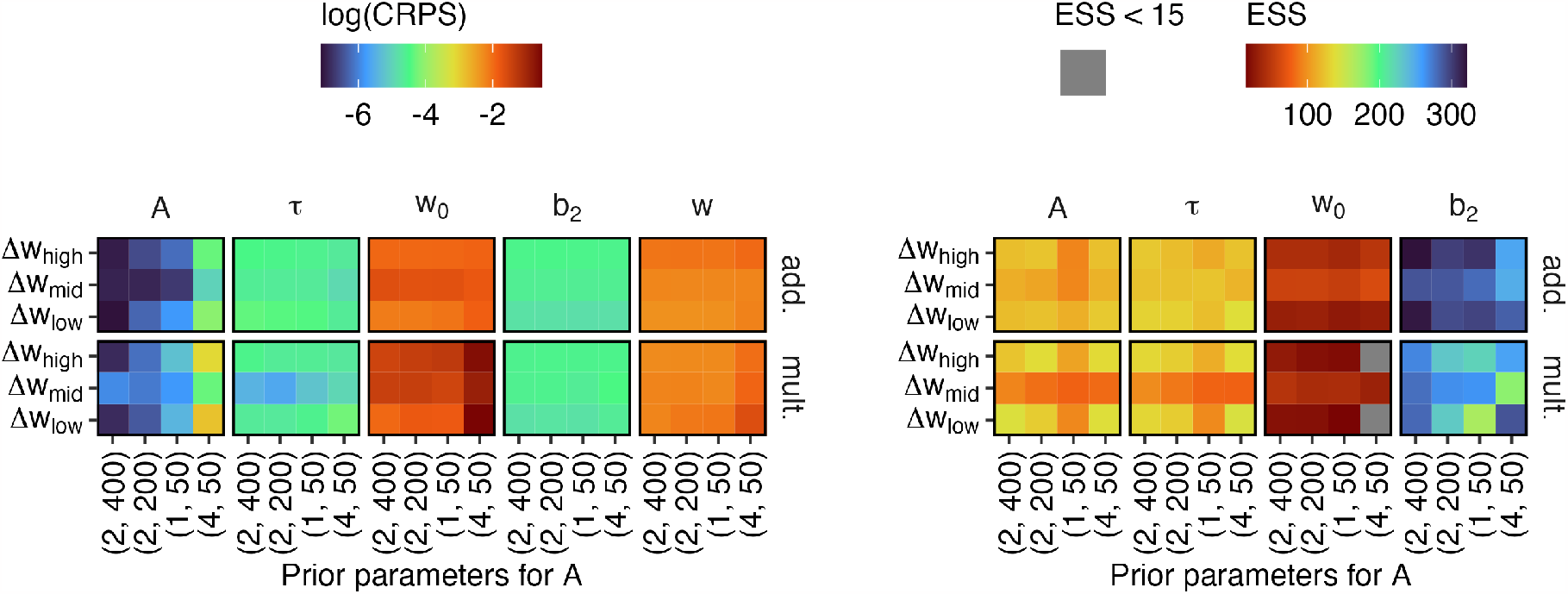
Performance when different prior parameters for the Gamma distribution on *A* are used. Sorted by increasing prior mean (*x*-axis) and for the different groups of weight trajectories (*y*-axis). log(CRPS) for the model parameters and weight trajectory (left) and ESS for the model parameters (right), as the mean over the 15, 53 and 32 model fits for each weight trajectory group (Δ*w*_low_, Δ*w*_mid_ and Δ*w*_high_) and prior. *w* represents the whole weight trajectory. We compute the average of the CRPS, and use the logarithm when visualizing the results. Inference for 5 multiplicative models with a Gamma(1, 50)-prior for *A* diverged.

**Fig. 7.**
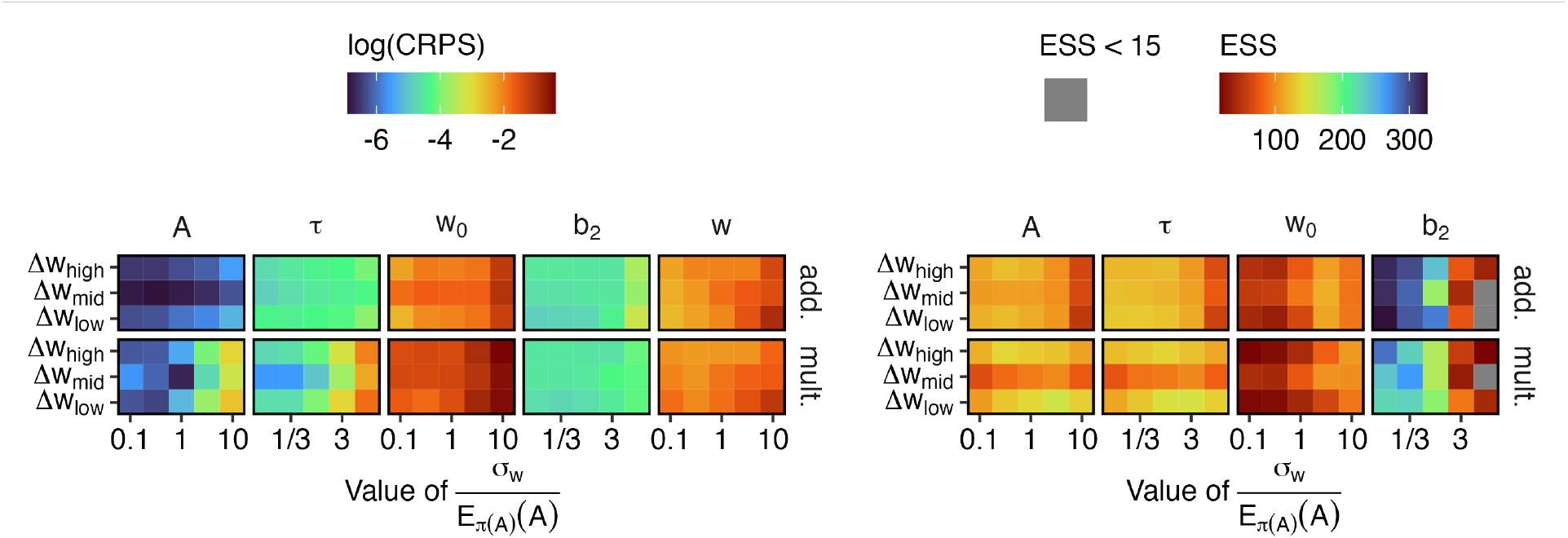
Performance when different prior parameters for the Gamma distribution on *τ* are used. Sorted by increasing prior mean (*x*-axis) and for the different groups of weight trajectories (*y*-axis). log(CRPS) for the model parameters and weight trajectory (left) and ESS for the model parameters (right), as the mean over the 15, 53 and 32 model fits for each weight trajectory group (Δ*w*_low_, Δ*w*_mid_ and Δ*w*_high_) and prior. *w* represents the whole weight trajectory. We compute the average of the CRPS, and use the logarithm when visualizing the results. Inference for 2 multiplicative models with a Gamma(1, 100)-prior for *τ* and 1 multiplicative model with a Gamma(5, 100)-prior for *τ* diverged.

### C.1.4 Value of weight trajectory noise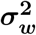

We studied how influential the fixed value of *σ*_*w*_ is on the inference (Fig. 8). A too high value leads to poor performance, especially with the multiplicative learning rule, while too low values will remove the randomness of the weight trajectory. Lower values perform better. We choose *σ*_*w*_ = *E*_*π*(*A*)_(*A*) *·* 1*/*3, since the random walk is assumed to be Gaussian, and 3 standard deviations covers more than 99.7% of the distribution. This means that in the prior model, the noise is almost never larger than the contribution of *A*, and the trend is mainly driven by the learning rule.

**Fig. 8.**
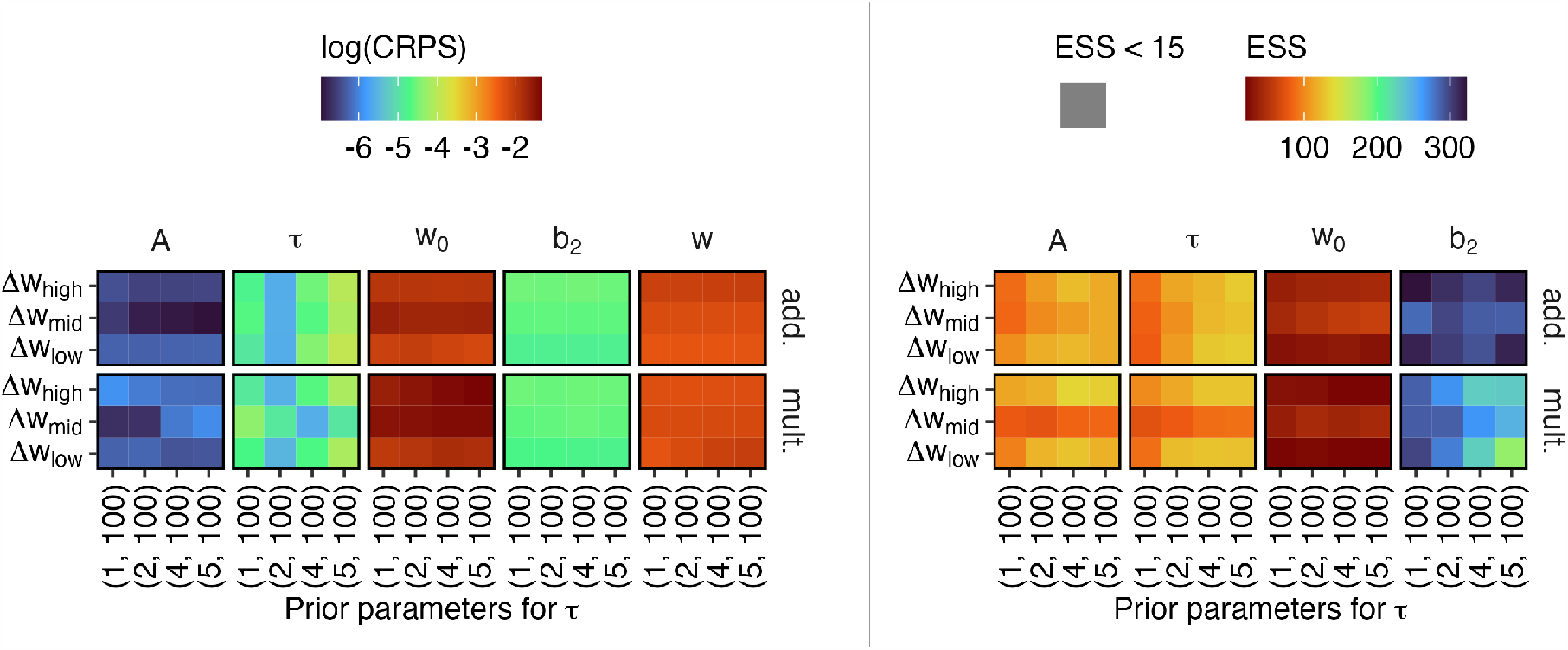
Performance when different values of the fixed noise parameter *σ*_*w*_ . Sorted by increasing value (*x*-axis) and for the different groups of weight trajectories (*y*-axis). The prior mean *E*_*π*(*A*)_(*A*) is 2*/*200. log(CRPS) for the model parameters and weight trajectory (left) and ESS for the model parameters (right), as the mean over the 15, 53 and 32 model fits for each weight trajectory group (Δ*w*_low_, Δ*w*_mid_ and Δ*w*_high_) and value of the noise. *w* represents the whole weight trajectory. We compute the average of the CRPS, and use the logarithm when visualizing the results. Inference for 4 multiplicative models with *σ*_*w*_ fixed to 0.1 *· E*_*π*(*A*)_(*A*) diverged.

### C.1.5 Different lag values

The correct algorithm lag setting (equal to the lag in the simulated data) performs better than using the wrong setting (different than the lag in the data) (Fig. 9). This is as expected and intended. Note that we have simulated 100 new datasets with each lag (1, 3, 5), in total 300 datasets, with the same values for *A, τ, w*_0_, *b*_1_, *b*_2_, *σ*_*w*_, time series length and learning rule as before.

**Fig. 9.**
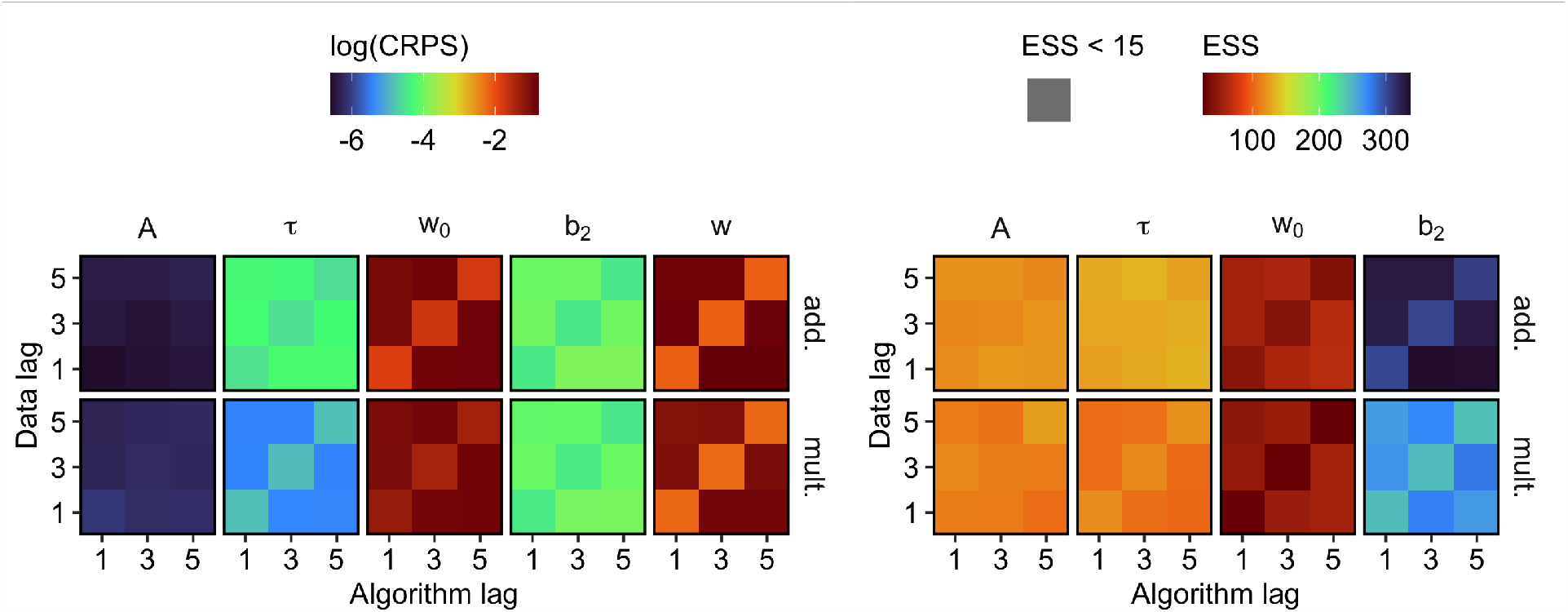
Performance for different lag values. Different lags used in the algorithm (*x*-axis) and simulated data (*y*-axis) (all combinations of lag 1, 3 and 5). log(CRPS) for the model parameters and weight trajectory (left) and ESS for the model parameters (right), as the mean over 100 simulated datasets for each scenario (lag value in data) and algorithm setting (lag value used by the algorithm). *w* represents the whole weight trajectory. We compute the average of the CRPS, and use the logarithm when visualizing the results. None of the model fits diverged.

## C.2 Model settings and model priors

We choose algorithm settings (Table 2) and model priors (Fig. 10) based on the simulated case study. Every *U* = 100 iteration, we adapt the variance of the proposal distributions according to the procedure of (Haario et al., 1999), using the newest *H* = 100 samples. The proposal standard deviation of *w*_0_ is restricted to be minimum 0.1 to ensure satisfactory mixing.

**Fig. 10.**
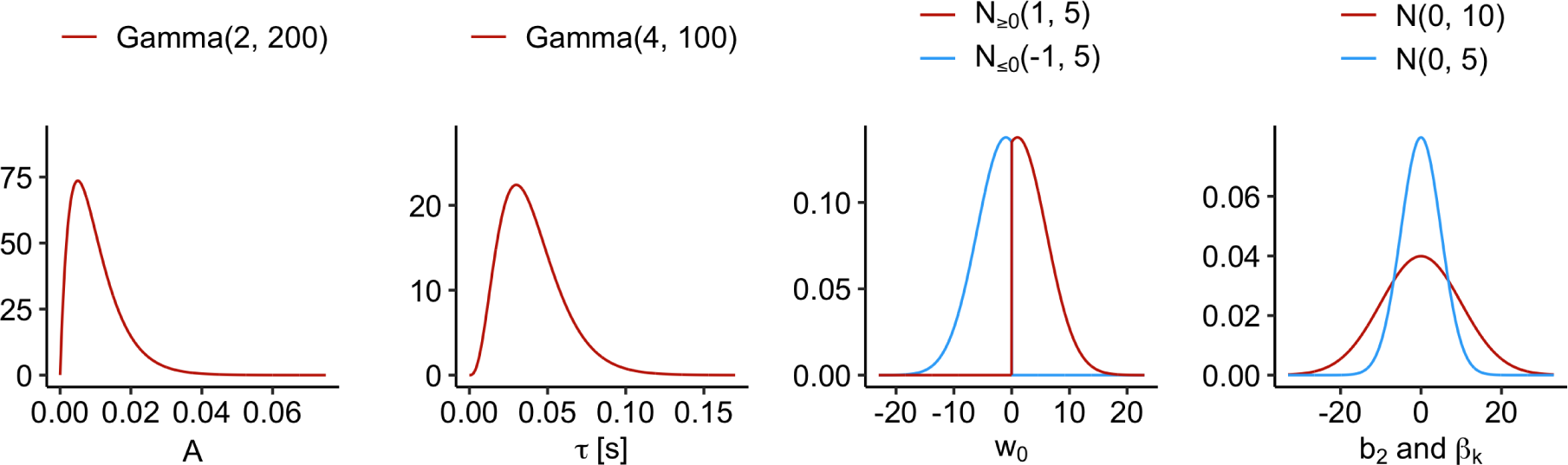
The prior distributions used for the model parameters in the inference. *b*_2_∼*𝒩* (0, 10^2^) and *β*_*k*_∼*𝒩* (0, 5^2^), and are displayed in the same graph for comparison (right).

## C.3 Model performance in various scenarios

We investigated how the model with the chosen settings (Table 2) and priors (Fig. 10) with both additive and multiplicative learning rule performs in various scenarios.

We varied the values of the true (simulated) *A, τ, w*_0_, *σ*_*w*_, length of time series, *b*_1_ and *b*_2_, covariate inclusion and which learning rule is used, when generating new datasets. We simulated 10 datasets for each setting. The covariate was simulated as a sine function with period 6.28 seconds and repeated for the length of the time series. We evaluated the performance using CRPS and ESS of the model parameters (Fig. 11 and Fig. 12).

**Fig. 11.**
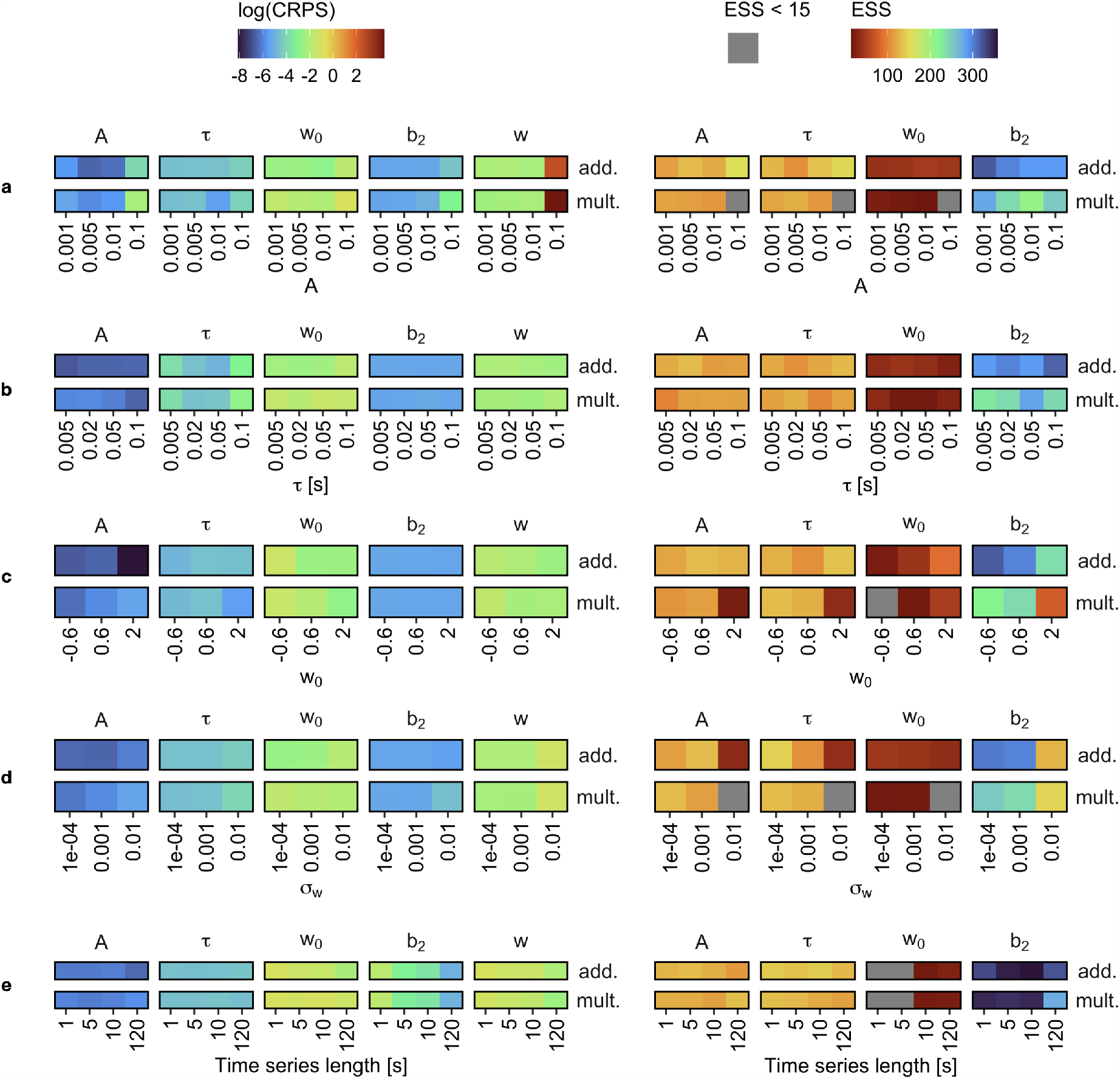
Algorithm performance in different scenarios. log(CRPS) for the model parameters and weight trajectory (left) and ESS for the model parameters (right), as the median over 10 simulated datasets. Different parameters/effects of true (simulated) dataset. We use the chosen settings (Table 2) and fit models with both additive and multiplicative learning rules (corresponding to rows). *w* represents the whole weight trajectory. We use the logarithm of the CRPS when visualizing the results. **a)** Different values of *A* (learning rule parameter). Inference for 1 multiplicative model with *A* = 0.1 diverged. **b)** Different values of *τ* (learning rule parameter). **c)** Different values of *w*_0_ (initial value of weight trajectory). **d)** Different values of *σ*_*w*_ (standard deviation of weight trajectory noise). **e)** Different length of the time series.

**Fig. 12.**
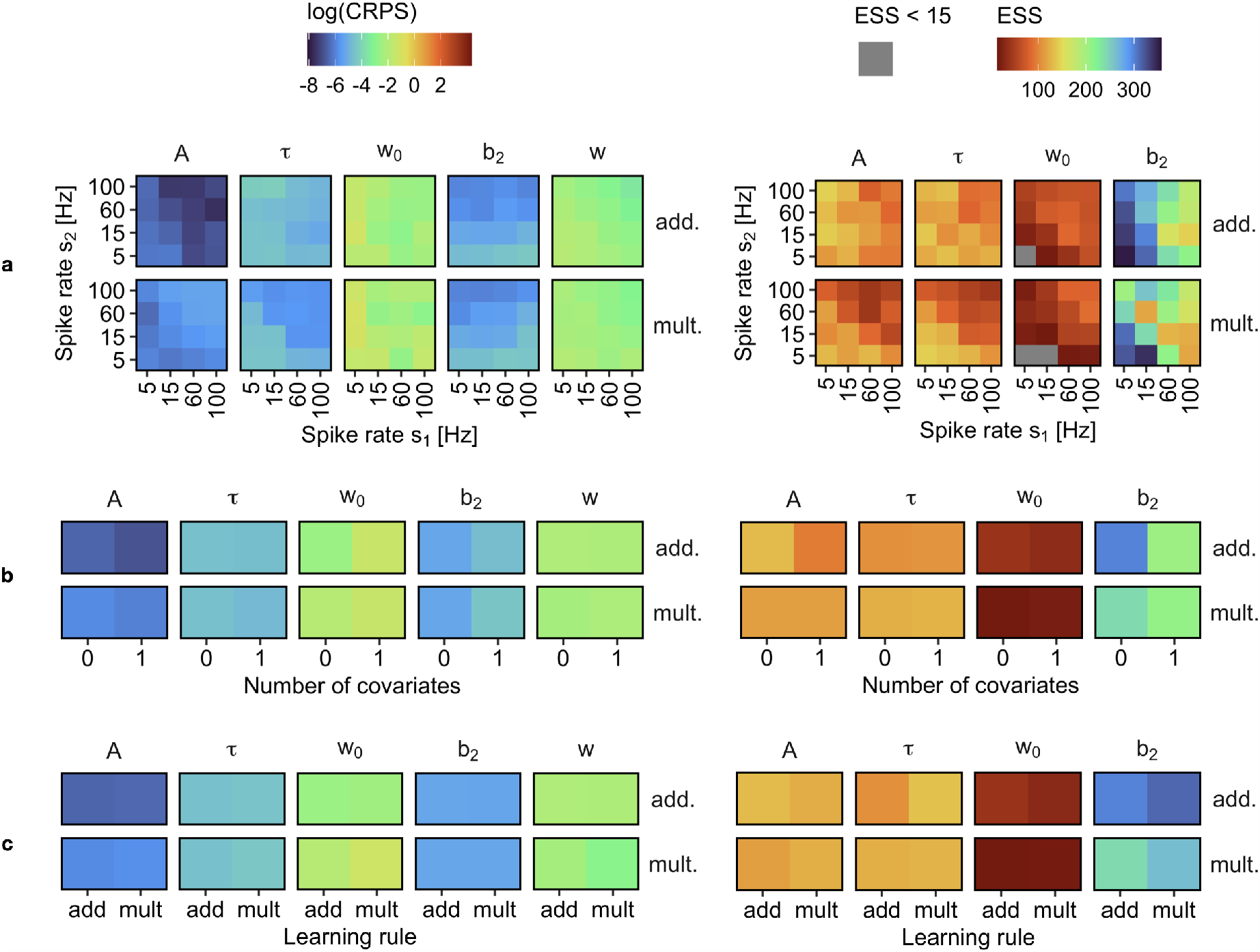
Algorithm performance in different scenarios. log(CRPS) for the model parameters and weight trajectory (left) and ESS for the model parameters (right), as the median over 10 simulated datasets. Different parameters/effects of true (simulated) dataset. We use the chosen settings (Table 2) and fit models with both additive and multiplicative learning rules (corresponding to rows). *w* represents the whole weight trajectory. We use the logarithm of the CRPS when visualizing the results. **a)** Different values of spike rate for both neurons 1 and 2 (i.e., different values of *b*_1_ and *b*_2_). Inference for 1 multiplicative model with spike rate of *s*_1_ equal to 100 and spike rate of *s*_2_ equal to 60 diverged. **b)** Without (0) and with (1) one covariate. The covariate is simulated as a sine function with period 6.28 seconds and repeated. We use *β*_1_ = 0.5. **c)** Additive and multiplicative learning rules for the simulated data.

A good performance means low CRPS and high ESS. Bad performance means high CRPS and/or low ESS. The additive model is in general performing equally good as or better than the multiplicative model, but the differences are bigger between normal and more extreme scenarios than between the different learning rules.

As long as the value of *A* is reasonable (less than 0.1), the model performs steady and well (Fig. 11**a**). The model performs slightly better for intermediate values of *τ* than for very high or very low values (Fig. 11**b**). The additive model is better at estimating *A* for large initial values of weight trajectory, *w*_0_, and the ESS for *w*_0_ is higher for higher values of true *w*_0_, but otherwise not much changes (Fig. 11**c**). For small values of *σ*_*w*_, the model performs well, and for *σ*_*w*_ = 0.01 the model performs worse with both additive and multiplicative learning rules (Fig. 11**d**). The model performs better for longer time series (Fig. 11**e**). Higher spike rates of both *s*_1_ and *s*_2_ leads to better model performance, but high spike rates of only one of them yields worse performance (Fig. 12**a**). Inclusion of covariates improves estimate accuracy of the learning rule parameters slightly, but leads to lower ESS (Fig. 12**b**). The additive and multiplicative models are both able to recover the true (simulated) values for data simulated with both learning rules (Fig. 12**c**).

