## Supplementary materials for "Bayesian inference of spike-timing dependent plasticity learning rules from single neuron recordings in humans"

### Supplementary figures

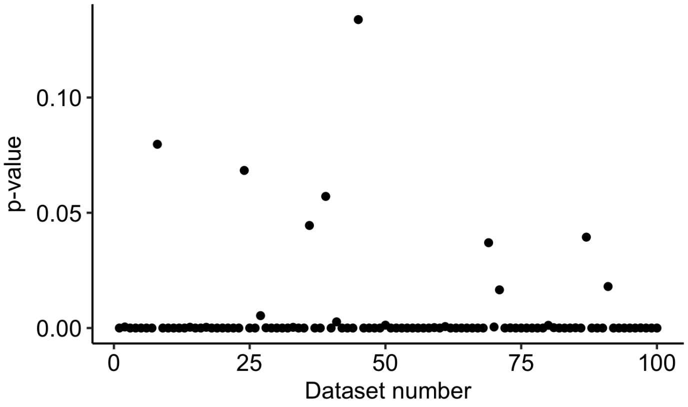


**Fig. S1**: Tail probabilities from temporal correlation investigation on simulated data. Tail probabilities for excitatory connections at time lag 1 from the temporal correlation between spike trains from the 100 simulated neuron pairs. The true time lag is equal to 1 time bin. The tail probability for excitatory connections is smaller than 0.025 at the correct lag in 93 of the 100 datasets.

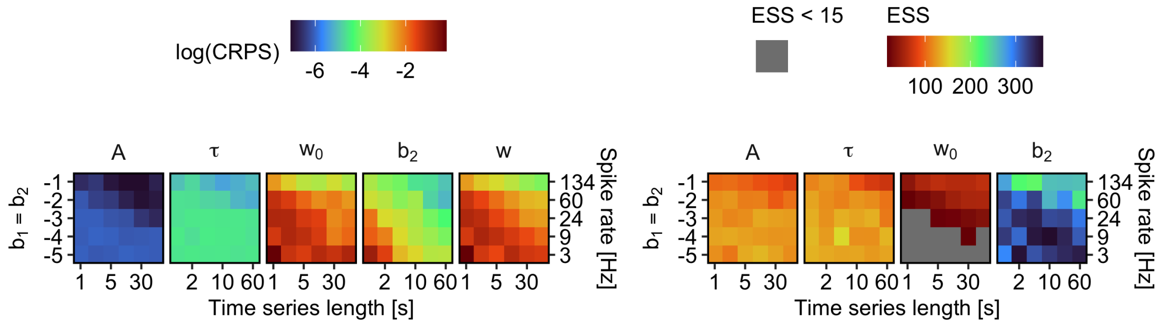


**Fig. S2**: CRPS and ESS for all parameters from scenarios with increasing values of $b_1 = b_2$ and time series length $T$. log(CRPS) for the model parameters and weight trajectory (left) and ESS for the model parameters (right), as the median over 10 simulated datasets for each combination of $b_1 = b_2$ and $T$. Additive learning rule only. $w$ represents the whole weight trajectory. We use the logarithm of the CRPS when visualizing the results. For $w_0$, we get median effective samples less than 15 for nearly half of the scenarios (combination of low $T$ and low $b_1 = b_2$). In longer time series with spike rates above 24 Hz, we get enough effective samples. No model fits crashed (stopped due to numerical errors) during the inference, and no model fits were removed even when the ESS was lower than 15. Longer datasets and higher spike rates give a lower (better) CRPS.

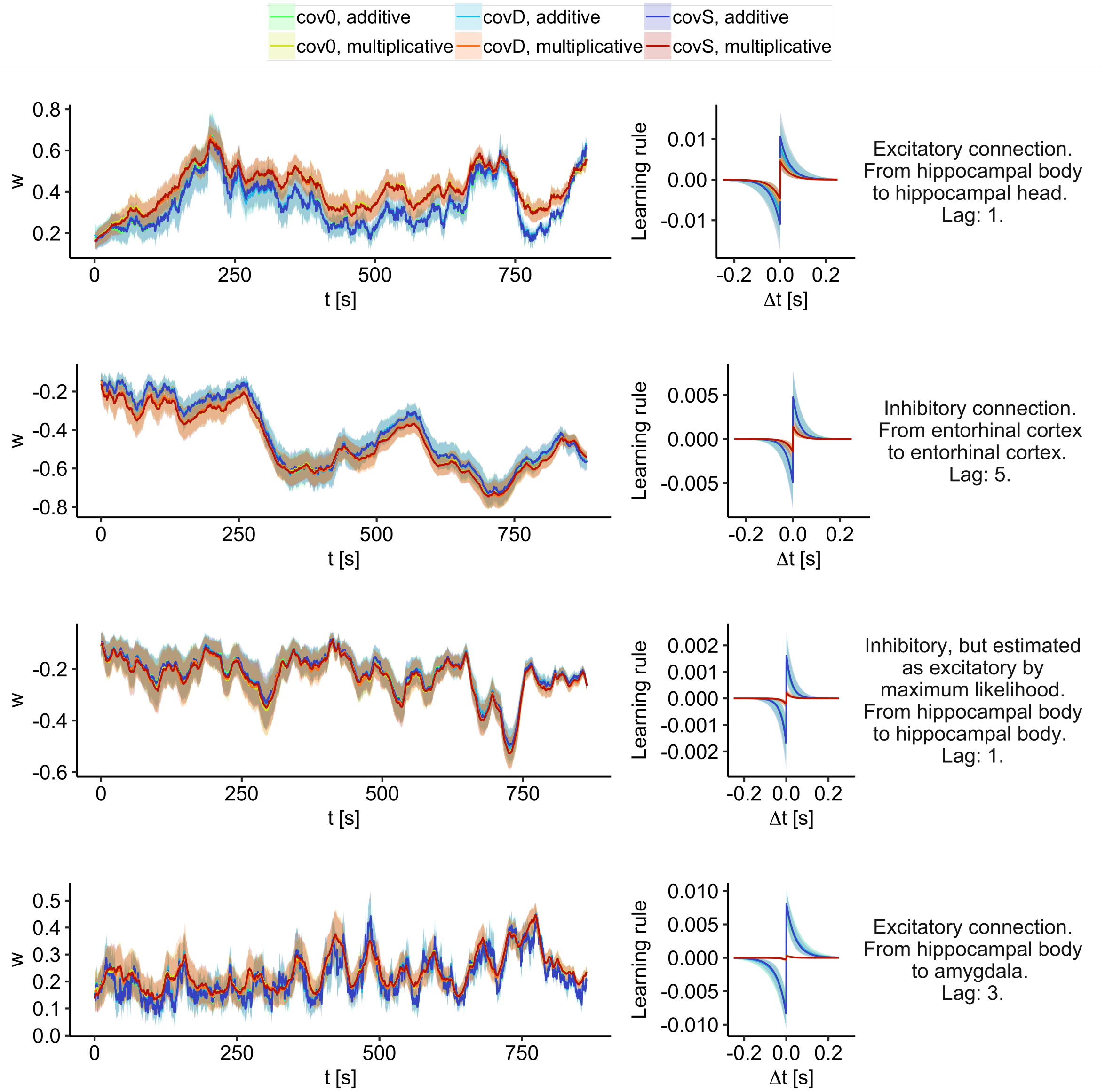


**Fig. S3**: Posterior weight trajectories and learning rules for both learning rule types and all covariate sets. Posterior weight trajectories (left) and corresponding learning rules (right) from four neuron pairs (one in each row, different subjects), for the two learning rules and three sets of covariates. Posterior weight trajectories and learning rules inferred using different covariates are almost perfectly overlaid, for the additive and multiplicative rule respectively. Connection types, brain areas and lags are indicated in the figure. The weight trajectories are displayed with posterior mean $\pm 1$ standard deviation. The learning rules are based on posterior medians and corresponding uncertainties on lower and upper quartiles of $A$ and $\tau$. Note that the interpretation of the additive and multiplicative learning rules are slightly different.

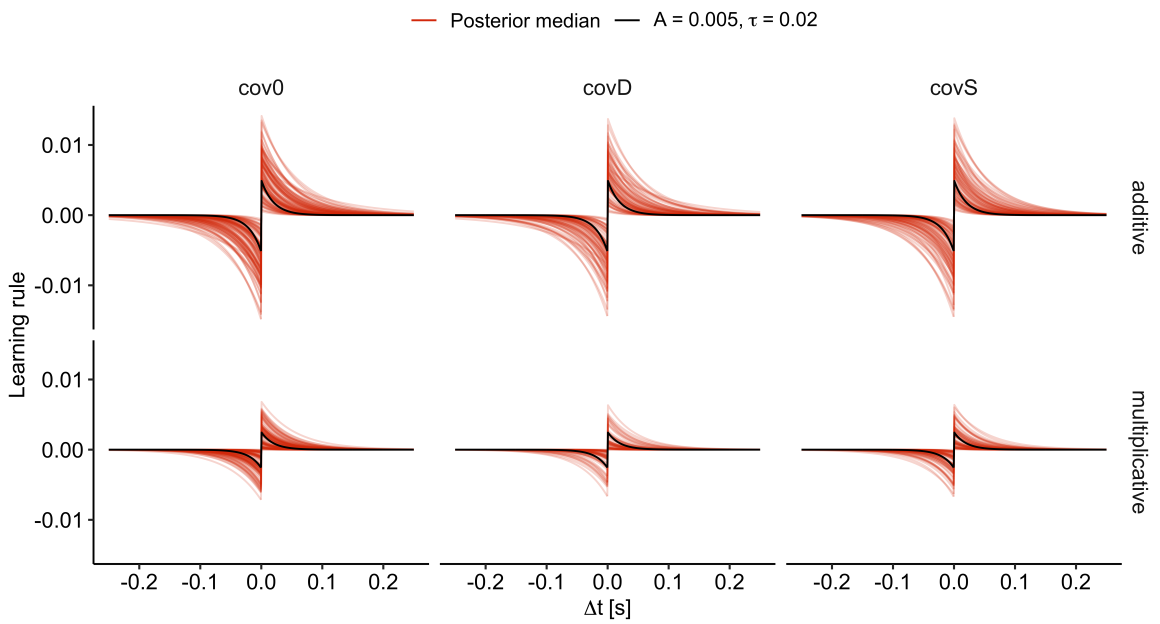


**Fig. S4**: Posterior learning rules for all model fits. Posterior median of $A$ and $\tau$ are used to visualize the learning rule for the model with additive (top) and multiplicative (bottom) learning rules, without covariates (cov0, left), with covariates describing the difficulty level of each trial (covD, middle) and covariates describing the stage of the trial (covS, right). We set $w_t = 0.5$ for all $t$ for the multiplicative learning rule. This is plotted with an additive learning rule with parameters $A = 0.005$ and $\tau = 0.02$ (top) and multiplicative learning rule with $A = 0.005$, $\tau = 0.02$ and $w_t = 0.5$. Only learning rules from converged model fits are included. Note that the $y$-axis vary between panels.

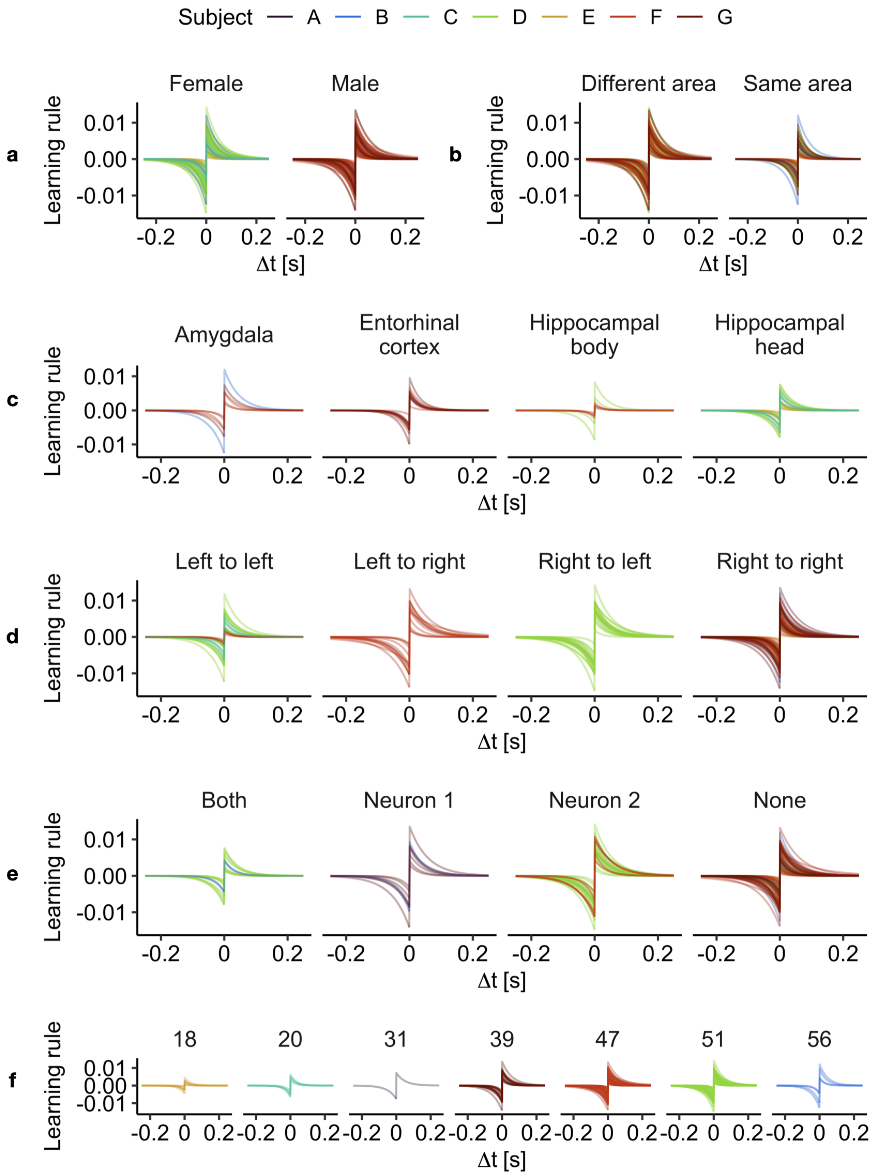


**Fig. S5**: Sorted learning rules, for the model with additive learning rule and no covariates. Posterior median of $A$ and $\tau$ are used to visualize the learning rule for the additive model without covariates (cov0). We sort them using different information about the data, that is not included in the inference. **a)** Gender of subject (male/female). **b)** Both neurons in a pair in the same or in two different areas. **c)** Of the neurons in the same area, which area are they located in. **d)** Which side of the brain neurons are located in (neuron 1 to neuron 2). **e)** Neurons located in seizure onset zones (both, neuron 1 ($\boldsymbol{s}_1$), neuron 2 ($\boldsymbol{s}_2$), or none). **f)** Age of subject.
